# B12 promotes gut dysbiosis and an inflammatory microenvironment that potentiates *Tet2*-deficient hematopoiesis

**DOI:** 10.1101/2025.08.22.671600

**Authors:** PD Lyon, TE Leesang, JP Brabson, KTT Do, M Dalzell, LA Nivelo, MQ Lam, A Peci, B Fang, V Strippoli, AA Armand, PK Singh, S Roy, F Beckedorff, AV Villarino, L Cimmino

## Abstract

Recent studies have linked elevated vitamin B12 serum levels with the presence of clonal hematopoiesis (CH) and an increased risk of developing myeloid malignancy. High B12 supplementation increases serum levels, alters gut microbial composition, and reduces the production of short-chain fatty acids (SCFAs), which help maintain gut barrier function and mucosal integrity. *TET2* mutation is a frequent driver of CH that progresses in a positive feedback loop in response to microbial signals suggesting that B12 may influence CH via the gut microbiome. We evaluated the microenvironmental effects of B12 supplementation in a *Tet2*-deficient model of CH and found that B12 enhances myelopoiesis, heightens the responses of myeloid cells to bacterial stimuli, and increases the levels of circulating inflammatory cytokines. B12 supplementation also induced gut dysbiosis and reduced the levels of SCFA-producing bacteria in both wild-type and *Tet2*-deficient mice. Importantly, the effects of excess B12 were reversible upon oral supplementation with the SCFA butyrate. These findings suggest that B12 may promote CH progression by disrupting microbiome-derived SCFA metabolism, highlighting a potential therapeutic role for SCFA supplementation in mitigating CH.

## Introduction

Gut dysbiosis has been implicated in the accelerated expansion of pre-malignant hematopoietic cells that harbor blood cancer-causing mutations, a condition known as clonal hematopoiesis (CH) (*1–3*). One of the most frequently mutated genes in CH and myeloid malignancies is *Ten-Eleven Translocation-2* (*TET2*), an epigenetic enzyme that regulates DNA demethylation and gene activation associated with myeloid cell differentiation. A hallmark of *TET2* loss of function is the aberrant self-renewal and expansion of myeloid-biased hematopoietic stem and progenitor cells (HSPCs), increased circulating myeloid cells, and elevated serum levels of pro-inflammatory cytokines (*4, 5*). *Tet2* deficiency also induces gut inflammation and increased intestinal permeability, resulting in bacteremia and a chronic pro-inflammatory state that accelerates pre-malignant HSPC and myeloid cell expansion in a positive feedback loop (*6, 7*). A germ-free environment reduces the expansion of *Tet2*-deficient cells in murine models (*6, 8*), highlighting the importance of the microbiome in the progression of pre-malignant hematopoiesis. Dietary changes rapidly alter the human gut microbiome (*9*) and the type of food a person consumes can have a significant impact on gut microbial diversity, and intestinal permeability that can lead to bacterial dissemination and systemic inflammation (*10, 11*). Short-chain fatty acids (SCFAs) such as propionate and butyrate are produced from intestinal bacteria (*12, 13*) and used directly by intestinal epithelial cells to strengthen the mucosal barrier (*14, 15*). One-carbon (1C) metabolism is a key regulator of SCFA production in the gut whereby dietary restriction of methionine has been shown to increase SCFA levels and reduce intestinal permeability and inflammation (*16, 17*). Increased B12 supplementation, the cofactor of methionine synthase that regulates methionine recycling, can also influence the gut microbiome, causing a reduction in microbial diversity and composition (*18, 19*). B12 is co-enriched with methionine in animal-derived foods, the consumption of which has been linked to CH prevalence (*20*). We recently reported that higher serum B12 levels are significantly associated with CH (*21*). This finding is consistent with other large population studies in Europe and China that reported an association between high B12 serum levels and increased risk of all-cause mortality, including heart disease, solid cancers, and myeloid malignancies (*22–26*). Given the association between B12 and CH and the inverse correlation between 1C metabolism, gut mucosal integrity, and SCFA production, we sought to determine whether B12 could affect the pre-malignant expansion of *Tet2*-deficient hematopoietic cells via modulation of the microbiome.

Here we report that high B12 supplementation exacerbates *Tet2*-deficient hematopoiesis, in a murine model of CH, resulting in the expansion of pre-malignant myeloid primed HSPCs, and elevated levels of circulating inflammatory cytokines. Fecal shotgun metagenomic sequencing analysis revealed that the changes caused by B12 were associated with a decrease in SCFA-producing bacteria in the gut. Furthermore, we find that B12 causes myeloid lineage cells with *Tet2* deficiency to respond more strongly to LPS stimulation by producing higher levels of *Il1b* and alarmins such as *S100a9*. Single-cell RNA sequencing of splenic myeloid cells and bone marrow cells further confirmed that high B12 supplementation promotes myeloid-biased differentiation and pro-inflammatory gene expression. Moreover, we show that SCFA supplementation in drinking water is sufficient to counteract the effects caused by excess B12, leading to a suppression of *Tet2*-deficient myeloid and HSPC expansion, reduced pro-inflammatory cytokine production, and a reversal of B12-induced gene expression changes. These findings highlight how interventions that target nutritional imbalances and gut microbial metabolism may aid in the prevention of inflammation and myeloid-biased hematopoiesis driven by *TET2* deficiency.

## Results

### High B12 treatment promotes a myeloid bias that exacerbates *Tet2*-deficient clonal hematopoiesis

To explore how B12 could impact normal and premalignant hematopoiesis, we performed competitive BM reconstitution assays with CD45.2^+^*Tet2*^+/+^ and *Tet2^+/−^* cells mixed 50:50 with CD45.1^+^ support cells (modeling a 25% VAF frequency in *TET2*) transplanted into lethally irradiated CD45.1^+^ recipient mice (Fig.1A). High B12 was supplemented in mice on a normal diet from 1-month post-transplant via intraperitoneal (IP) injection of 0.29µg in PBS, equivalent to a 1mg B12 dose in a 70kg human, twice per week. This dosing regimen mimics treatment provided to patients receiving high parenteral B12 supplementation in clinical practice (*27, 28*) and was based on the observation that a single dose of B12 IP was sufficient to raise B12 serum levels ∼1.5-fold above baseline for up to 72 hours (Fig.S1A). Low B12 and standard B12 supplementation were modeled by providing low or normal B12 dietary supplementation in rodent chow.

**Figure 1:**
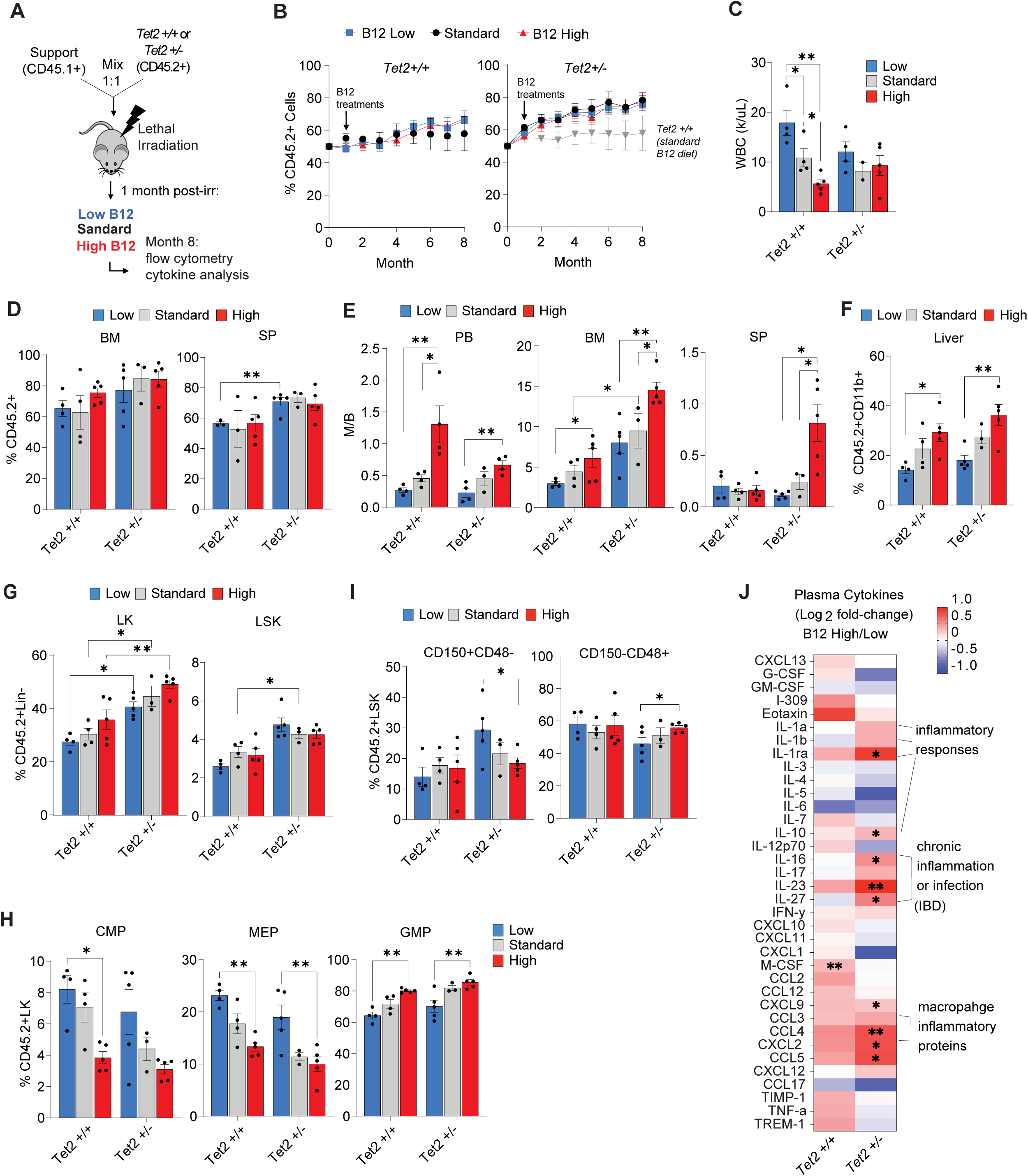
High B12 supplementation exacerbates *Tet2*-deficient myelopoiesis in mice. (A) Schematic of CD45.2^+^ *Tet2^+/+^* and *Tet2^+/−^* competitive bone marrow (BM) reconstitution (mixed 1:1) with CD45.1^+^ wild-type BM cells transplanted into congenic mice. Altered B12 supplementation was initiated from 1-month post-transplant and mice were monitored for 8 months prior to sacrifice. (B) Frequency of peripheral blood CD45.2^+^ cells over 8 months post-transplant in *Tet2*^+/+^ and *Tet2*^+/−^ reconstituted mice with altered B12 supplementation. (C-J) Hematopoietic phenotypes of competitively transplanted mice at 8 months post-transplant. (C) Total WBC counts of cohorts (k/µL). (D) Frequency of CD45.2^+^ cells in peripheral blood (PB), bone marrow (BM), and spleen (SP). (E) Myeloid-to-B-cell (M/B) ratio measured by flow cytometry of CD11b^+^ vs B220^+^ cells in the peripheral blood (PB), bone marrow (BM), and spleen (SP). (F) Frequency of CD45.2^+^CD11b^+^ cells in the liver. (G) Frequency of CD45.2^+^ Lineage negative (Lin-) cKit^+^ (LK) cells, and lineage negative cKit^+^ Sca1^+^ (LSK) cells. (H) Frequency of common myeloid progenitor (CMP), megakaryocyte and erythroid progenitor (MEP), and granulocyte and macrophage progenitor (GMP) cells in the CD45.2^+^ LK compartment. (I) Frequency of LSK cells that are CD150^+^CD48^-^ (HSCs) and CD150^-^CD48^+^ (myeloid primed multipotent progenitors) within the BM compartment. (J) Relative plasma cytokine levels in B12 high-treated mice compared to B12 low in both *Tet2^+/+^* and *Tet2^+/−^* cohorts. Panels show mean and STD of n = 4-5 mice per group, *p < 0.05, **p < 0.01, ***p < 0.001.

We observed that while CD45.2^+^*Tet2*^+/−^ hematopoietic cells showed a competitive advantage in the peripheral blood (PB) compared to *Tet2^+^*^/+^ as previously reported (*29, 30*), there was no effect of altered B12 supplementation (Fig.1B). Total white blood cells (WBCs) showed variability in response to altered levels in the context of *Tet2*^+/+^ hematopoiesis, red blood cells (RBCs), hemoglobin (Hgb), and mean corpuscular volume (MCV) were all within normal ranges for both genotypes (Fig.1C, Fig.S1B). Despite a significant increase in the frequency of *Tet2*^+/−^ compared to *Tet2*^+/+^ (CD45.2^+^) donor cells in the spleen (SP) of low B12 supplemented mice, the SP and bone marrow (BM) reconstitution frequencies for each genotype were unaltered by the different levels of B12 in mice sacrificed after 7 months of treatment (8 months post-transplant) (Fig.1D). High B12 supplementation did however significantly increase the myeloid-to-B-cell (M/B) ratio of both *Tet2*^+/+^ and *Tet2*^+/−^ cells in PB, BM and SP (Fig.1E, Fig.S1C), with a strong myeloid bias observed in the SP specifically in the context of *Tet2*-deficient hematopoiesis. The frequency of CD45.2^+^ *Tet2*^+/+^ and *Tet2*^+/−^ myeloid cells in the liver (Fig.1F) also increased in response to high B12 supplementation. In the BM compartment, lineage negative, cKit^+^ (LK) cells showed a trend toward increasing frequency by high B12 supplementation but was not statistically significant, whereas total lineage negative cKit^+^Sca1^+^ (LSK) cells were unchanged by altered B12 treatment (Fig.1G). In the LK compartment, the increased frequency of CD45.2^+^ cells was associated with a significant expansion of granulocyte and myeloid progenitors (GMPs) in both *Tet2*^+/+^ and *Tet2*^+/−^ donor cells (Fig.1H). However, within the LSK compartment, a significant reduction in HSCs (CD150^+^CD48^-^) and an increase in myeloid-primed multipotent progenitors (CD150^-^CD48^+^) was observed only for CD45.2^+^*Tet2*^+/−^ cells from high B12 treated reconstituted mice (Fig.1I). These experiments show that increased B12 supplementation *in vivo* causes a myeloid cell lineage differentiation bias that cooperates to enhance *Tet2* deficient pre-malignant myelopoiesis.

We next sought to determine the impact of B12 on systemic inflammatory signaling given *Tet2*-deficient CH is associated with the development and differentiation of pro-inflammatory myeloid cells (*7, 31*). Serum cytokine analysis showed that high B12 supplementation increased circulating inflammatory mediators including IL-1α, IL-1β, IL-RA, IL-23, IL-27, CCL-4/5, and CXCL-2 (Fig.1J) in both *Tet2*^+/+^ and *Tet2^+/−^*reconstituted mice, but more strongly in the latter, indicating a cooperation between B12 levels and *Tet2*-deficiency. Many of the serum cytokines found to be increased with high B12 supplementation have been reported in patients with CHIP and myeloid neoplasms including IL-1β (CHIP, myeloproliferative neoplasm-MPN), IL-1RA (MDS, AML, MPN), CXCL-2 – an orthologue to human IL-8 (CHIP, MDS, AML, MPN), CCL-4 (MDS, MPN), CXCL-9 (MDS, MPN), and IL-10 (AML, MPN)(*3, 32, 33*). We also tested whether high dietary consumption of B12 as opposed to IP injections could influence *Tet2*-deficient CH (Fig.S1D) and show that in BM reconstituted mice, after 12 months of dietary intervention, the same cellular phenotypes were induced (Fig.S1E-I), notably, a strong bias in the M/B cell ratio across all hematopoietic organs (Fig.S1G), and significantly increased LK and GMP cell frequencies (Fig.S1H-I). These data indicate that high B12 supplementation can increase circulating B12 serum levels and promote a *Tet2*-deficient myeloid differentiation bias and inflammatory microenvironment.

### B12 increases innate inflammatory gene expression and the expansion of *Tet2*-deficient pro-inflammatory myeloid cells

To understand how B12 influences *Tet2*-deficient myeloid cell phenotypes, we performed scRNA-sequencing of *Tet2*^+/−^ splenic CD45.2^+^CD11b^+^ cells from B12 low and high supplemented host mice. We observed an increase in neutrophils (cluster 0 and 6) but decrease in antigen-presenting memory-like CD11b^+^ B cells (cluster 1) according to Immgen cluster identification (*34*) (Fig.2A-B). While B12 treatment appeared to decrease the frequency of monocytes (cluster 5), pseudotime analysis suggested that this may be due to their enhanced differentiation into neutrophils (cluster 0) (Fig.2C).

**Figure 2:**
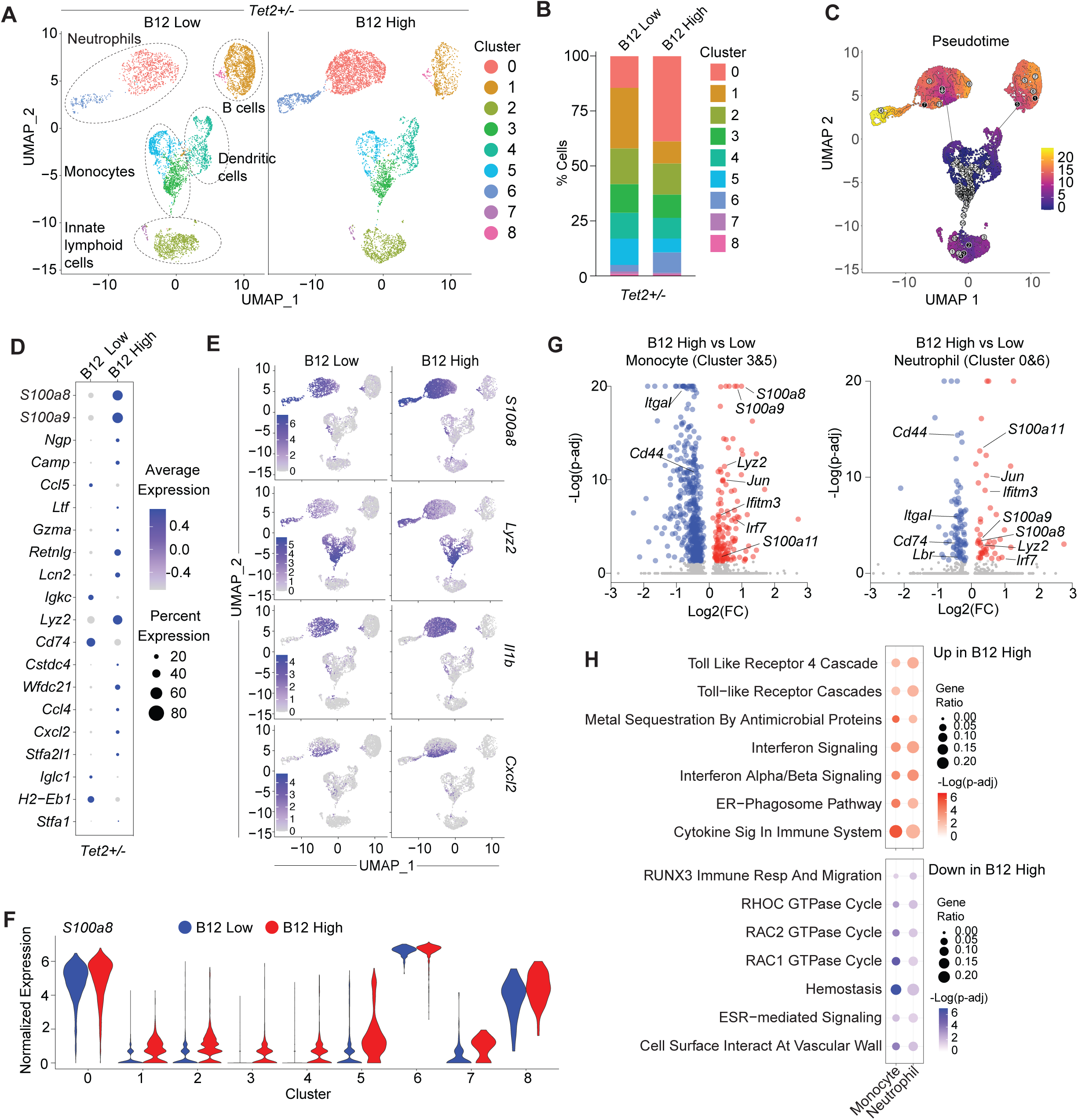
B12 increases innate inflammatory gene expression and expansion of *Tet2*-deficient pro-inflammatory myeloid cells. (A) UMAP plots of scRNAseq classified using Immgen cell references with clustifyr from CD45.2^+^CD11b^+^ sort-purified splenic cells of *Tet2^+/−^* BM reconstituted mice supplemented with low (left) and high (right) B12. (B) Relative frequency of cells in each cluster per B12 supplemented group. (C) Global pseudotime analysis of cells across both treatment groups. (D) Average gene expression of the top 20 cluster-defining markers in B12 high and B12 low supplemented conditions. (E) UMAP plots of relative expression of *S100a8*, *Lyz2, Il1b,* and *Cxcl2* split by treatment arm and scaled to the maximum value. (F) Violin plots of normalized expression of *S100a8* across all clusters. (G) Volcano plot of differentially expressed genes (DEGs) from Monocyte (Cluster 3&5) and Neutrophil (Clusters 0 and 6) cells with a significance cutoff of padj < 0.05 in blue (down in B12 high) or red (up in B12 high). (H) Enriched gene sets of DEGs in the Neutrophil and Monocyte clusters from Reactome Pathway Database. Significance cutoff of padj < 0.05.

Analysis of top globally– and cluster-variable genes in *Tet2*^+/−^ splenic CD11b+ cells showed that *S100 calcium binding protein A8* (*S100a8*) and *A9* (*S100a9*) were amongst the top genes upregulated upon high B12 supplementation across all major clusters (Fig.2D-F, Fig.S2A-B). *Lipocalin2* (*Lcn2*), a known mediator of neutrophilic inflammation that limits bacterial growth through iron sequestration (*35*), and *WAP four-disulfide core domain 21* (*Wfdc21*) that plays a role in Toll-like receptor 4 (TLR4) signaling (*36*) were also upregulated in high B12 supplemented mice (Fig.2D, Fig.S2A). In monocytes (cluster 5), we observed increased expression of Histocompatability 2 genes (*H2-Aa*, *H2-Ab1*, *H2-Eb1*) (Fig.S2A) which are homologs to the human MHCII genes (*37*) and more monocytes and/or neutrophils were found to be expressing pro-inflammatory cytokines such as *Interleukin 1 beta* (*Il1b*) along with *S100a8*, *Lyz2*, and *Cxcl2* upon high B12 supplementation (Fig.2E-F).

Differential gene expression (DEG) analysis in monocytes (cluster 3&5) and neutrophils (clusters 0&6) further confirmed a significant increase in expression of alarmins (*S100a8*, *S100a9, S100a11*), interferon response mediators (*Ifitm3*, *Irf7*, *Ifi27*), and other genes associated with innate inflammatory responses (*Lyz2*, *Jun*) in high B12 supplemented mice (Fig.2G) that overlapped between neutrophils and monocytes (Fig.S2C and Table S2). Pathway analysis of genes expressed in monocyte and neutrophil clusters showed enrichment of TLR4, interferon, and IL-17 signaling, while RHOC and RAC1/2 GTPase Cycle genes were depleted upon high B12 supplementation (Fig.2H, Fig.S2D-E). These data revealed that the increased frequency of myeloid cells in the spleen of high B12 supplemented mice exhibit upregulated expression of innate inflammatory genes known to play a role in the progression of pre-malignant hematopoiesis and myeloid malignancy (*6, 31*).

### High B12 supplementation causes gut dysbiosis and a reduction in fecal SCFA-producing bacteria

Increased B12 supplementation is known to influence the gut microbiome, causing a reduction in microbial diversity and composition (*18, 19*). *Tet2*-deficient hematopoiesis is fueled by gut microbial signaling (*6*), hence, we sought to determine whether B12 could affect the pre-malignant expansion of *Tet2*-deficient hematopoietic cells via modulation of the microbiome. We performed shotgun sequencing and metagenomic analysis of fecal pellets isolated at month 7 of B12 supplementation (8 months post-transplant) from *Tet2*^+/+^ and *Tet*2^+/−^ BM reconstituted mice (Fig.3A). We found that high B12 causes a significant decrease in α-diversity in the context of both genotypes according to the Shannon and Simpson’s Indices (Fig.3B), which are measurements of species richness and evenness within a sample (*38*). Analysis of the phyla showed an increase in Actinobacteria, while Firmicutes and Bacteroidetes decreased in frequency (Fig.3C). 1C metabolism has previously been shown to regulate gut bacterial SCFA production (*16, 17*) and at the genera level, SCFA-producing bacteria were significantly affected by high B12 supplementation, which caused a decrease in butyrate producers in both *Tet2*^+/+^ and *Tet2*^+/−^ hosts (Fig.3D). Linear discriminant analysis effect size (LEFSE) of bacterial species in high B12 supplemented cohorts showed depletion of commensal genera *Lactobacillus*, *Lachnospiraceae and Enterohabdus* (Fig.3E-F). Of these, a decreased level of *Lachnospiraceae* has been associated with an increased neutrophil-to-lymphocyte ratio in aged mice and *Enterorhabdus* genera are normally associated with healthy aging (decreased frailty index) (*39*). *Akkermansia muciniphila* was enriched in high B12 supplemented mice, and while this bacterial species is described as a protective probiotic for several diseases (*40*) its increased presence correlates with intestinal barrier damage and worsening of colitis-associated disease (*41, 42*). Finally, KEGG pathway metagenomic analyses showed reduced propanoate and butanoate metabolism in the high B12 supplemented mice, indicating a loss in the production of protective SCFAs (Fig.3G). These data show that B12 promotes a myeloid bias, gut dysbiosis, and decreased SCFA-producing bacteria.

**Figure 3:**
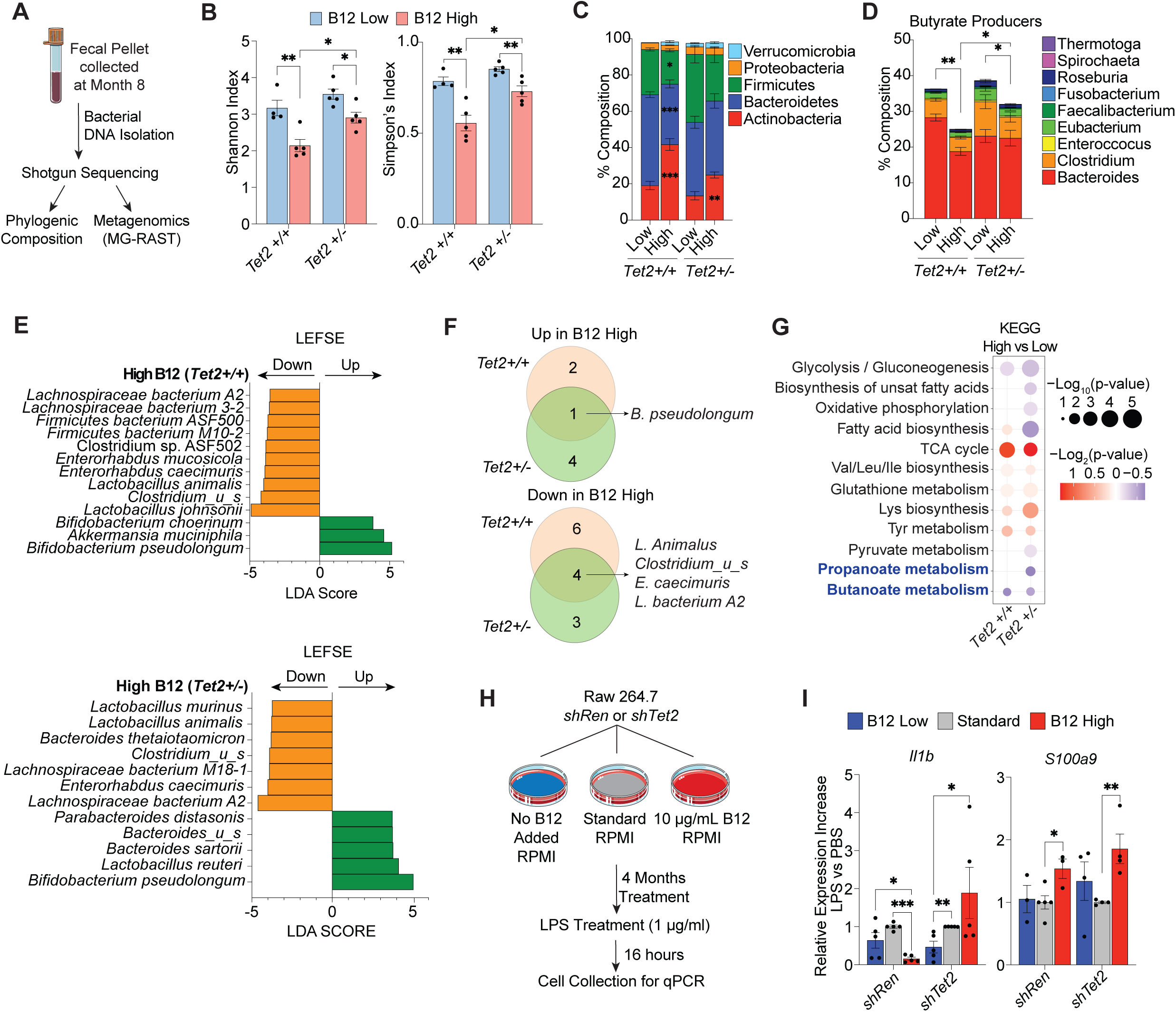
High B12 supplementation causes gut dysbiosis leading to a reduction in SCFA-producing bacteria. (A) Schematic of collection at month 8, bacterial DNA isolation, and shotgun metagenomic sequencing analysis of fecal pellets from mice (n= 4-5 mice per group) with competitive transplant of CD45.2^+^ *Tet2*^+/+^ and *Tet2*^+/−^ BM cells supplemented with low to high B12 (as outlined in Figure 1). (B) Alpha diversity levels measured by Shannon and Simpson’s indices. (C) Fecal microbiome composition at the phylum level averaged per group from B12 supplemented cohorts of *Tet2^+/+^*and *Tet2^+/−^* host mice. (D) Composition levels of butyrate producers at the genera level in the fecal microbiome of B12 supplemented *Tet2^+/+^* and *Tet2^+/−^* host mice. (E) Bar graphs of linear discriminant analysis (LDA) score calculated by linear discriminant analysis effect size (LEFSE) for species in the fecal microbiome. (F) Venn Diagrams of overlaps of enriched or depleted species in the high vs low B12-treated *Tet2^+/+^* and *Tet2^+/−^* competitive BM reconstituted mice. (G) Dot plot of enriched KEGG pathways from metagenomic analysis comparing the fecal microbiomes of respective low and high B12 supplemented cohorts using the MG-RAST pipeline. (H) Experimental schematic of *shRen* and *shTet2* Raw 264.7 mouse macrophage cells cultured in low, standard, and high B12 supplemented media. (I) Gene expression levels of *Il1b* and *S100a9* in Raw 264.7 cells after 16-hour stimulation with PBS or 1 µg/mL LPS. p-values = *p < 0.05, **p < 0.005, ***p < 0.0005.

*Tet2*-deficient hematopoiesis itself drives intestinal inflammation, bacterial dissemination, and an increase in circulating inflammatory mediators in a positive feedback loop that can promote the expansion of myeloid cells in mice under normal dietary conditions (*6*). We therefore assessed whether *Tet2*-deficient immune cells promote gut dysbiosis independently of B12 by comparing shotgun sequencing and SCFA producers of fecal pellets between mice with *Tet2^+/+^*or *Tet2^+/−^* competitive BM reconstitution (Fig.S3A-F). While *Tet2*-deficient hematopoiesis alone did not alter α-diversity in murine hosts (Fig.S3G), a significant increase in the frequency of Actinobacteria and a reduction in Bacteroidetes phyla was observed (Fig.S3H). At the genera level, a small but significant decrease in the frequency of propionate and butyrate SCFA-producing bacteria (Fig. S3I) and a depletion of *Lactobacilli* was also associated with *Tet2*^+/−^ compared to *Tet2*^+/+^ hematopoeisis (Fig.S3J) in addition to changes in metabolic KEGG pathway activity (Fig.S3K). These data show that *Tet2*-deficient hematopoiesis alone can deplete commensal anti-inflammatory bacteria such as SCFA producers.

B12 may directly influence myeloid cell inflammatory signaling independently of its effect on the microbiome. The elevation in circulating IL-1 signaling mediators observed upon high B12 supplementation may also lead to enhanced responsiveness to microbial signaling as previously described for IL-6 (*6*). We therefore tested the effect of low to high B12 supplementation in combination with microbial signaling via lipopolysaccharide (LPS) using RAW 264.7 myeloid mouse cells transduced with control *shRenilla* (*shRen*) or *shTet2* (Fig.3H). Upon LPS stimulation, RAW 267.4 cells cultured in the presence of high B12 exhibited increased expression of *Il1b* only in the context of *Tet2* knockdown, whereas alarmin gene *S100a9* expression was upregulated in both control and *Tet2* knockdown cells (Fig.3I). These data suggest that high B12 levels can directly enhance the expression of inflammatory mediators in response to bacterial stimuli, thereby exacerbating *Tet2*-deficient phenotypes through cell-intrinsic and microenvironmental effects.

### SCFA treatment ameliorates the myeloid bias and inflammatory microenvironment caused by high B12 supplementation

Treatment with SCFAs, such as butyrate, has been shown in murine models of AML to reduce intestinal permeability and slow disease progression (*43*). Whether SCFAs can also suppress the phenotypes of CH progression has not been described before. Based on the observation that high B12 supplementation leads to a depletion of butyrate-producing gut bacteria, we investigated whether exogenous butyrate supplementation via drinking water could offset the impact of high B12 on hematopoiesis in our competitive *Tet2*-deficient bone marrow transplantation model (Fig.4A). Similar to our previous results, we found that high B12 and butyrate treatment alone or in combination did not alter WBC, RBC, HgB or MCV levels (Fig.4B, Fig.S4A), nor the competitiveness of *Tet2*^+/−^ cells in the PB, SP or BM (Fig.4B). However, at the completion of 7 months treatment (8 months post-transplant), there was a significant increase in the M/B cell ratio induced by high B12 supplementation that was reversed by co-treatment with butyrate (Fig.4C, Fig.S4B). These data correlated with an overall reduction in B12-upregulated circulating inflammatory cytokines upon butyrate treatment (most significantly for IL-6 and CXCL-12) (Fig.4D-E). Analysis of the bone marrow also showed that butyrate suppressed the B12-induced expansion of LK and GMP cells, consistent with reversion of a myeloid lineage differentiation bias (Fig.4F-G). Support CD45.1^+^*Tet2*^+/+^ cells also displayed an increased M/B cell ratio in high B12 treated mice that was reversed by butyrate supplementation (Fig.S4C), further suggesting that B12 alters the microenvironment to influence myeloid differentiation that effects both pre-malignant and non-malignant hematopoietic cells, as shown in independent *Tet2*^+/+^ and *Tet2*^+/−^ hosts in Figure 1. We performed a FITC dextran oral gavage and translocation assay to measure intestinal permeability before sacrifice at 8 months post-transplant. Though not statistically significant, a notable trend was observed in which increased FITC-dextran translocation to the peripheral blood in high B12 treated mice (Fig.S4D) was reversed upon butyrate supplementation. Taken together, these data suggest that butyrate treatment is sufficient to block the pro-inflammatory signaling and myeloid bias caused by increased B12 supplementation that may exert is systemic effects via altered gut permeability.

**Figure 4:**
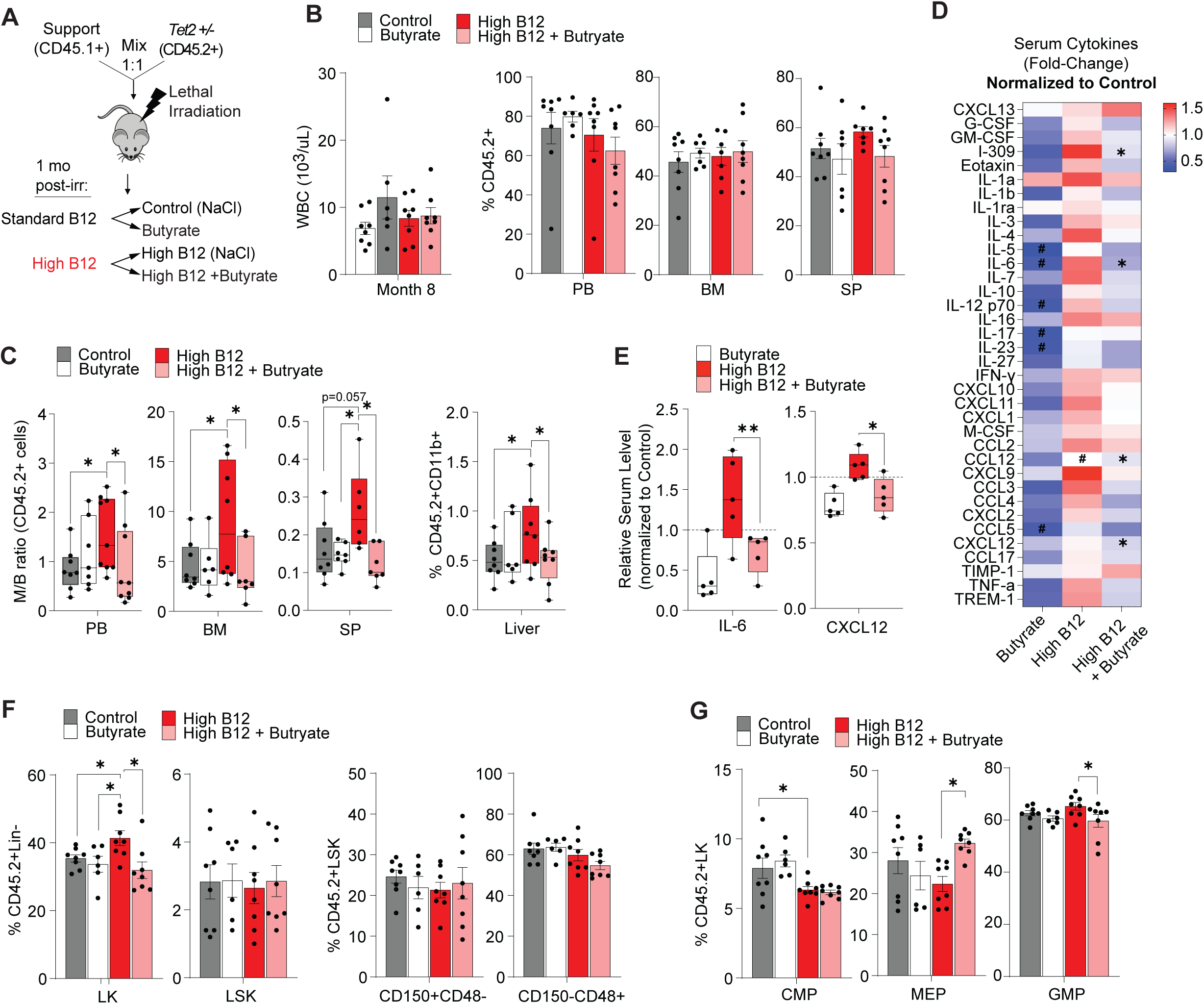
Butyrate treatment ameliorates the inflammatory microenvironment and enhanced myelopoiesis of *Tet2*-deficient cells caused by high B12 supplementation. (A) Schematic of CD45.2^+^ *Tet2^+/−^* competitive bone marrow (BM) reconstitution (mixed 1:1) with CD45.1^+^ wild-type BM cells transplanted into congenic mice. Altered B12 supplementation and oral butyrate treatment (as sodium butyrate provided in drinking water) was initiated from 1-month post-transplant and mice were monitored for 8 months prior to sacrifice. (B-G) Hematopoietic phenotypes of competitive BM transplanted mice 8 months post-transplant. (B) Total WBC counts of treatment cohorts (k/µL, left panel) and frequency of total CD45.2^+^ in peripheral blood (PB) bone marrow (BM) and spleen (SP) (right panels). (C) Myeloid-to-B-cell (M/B) ratio measured by flow cytometry of CD45.2^+^CD11b^+^ vs B220^+^ cells in the peripheral blood (PB), bone marrow (BM), and spleen (SP) and the frequency of CD45.2^+^CD11b^+^ cells in the liver of (right panel). (D) Relative serum cytokine levels in mice treated with butyrate, high B12, and high B12+butyrate (normalized to control mice). *p < 0.05 compared to High B12, #p<0.05 compared to Control. (E) Relative IL-6 and CXCL12 levels in the serum of butyrate, high B12, and high B12+butyrate *Tet2*^+/−^ host mice. (F) Frequency of CD45.2^+^ Lineage negative (Lin-) cKit^+^ (LK) cells, lineage negative cKit^+^ Sca1^+^ (LSK) cells and LSK subsets that are CD150^+^CD48^-^ (HSCs) and CD150^-^CD48^+^ (myeloid primed multipotent progenitors) within the BM compartment. (G) Frequency of common myeloid progenitor (CMP), megakaryocyte and erythroid progenitor (MEP) and granulocyte and macrophage progenitor (GMP) cells in the CD45.2^+^ LK compartment. Panels show mean and STD of n = 5-8 mice per group, *p < 0.05, **p < 0.005, unless otherwise stated above.

### SCFA treatment suppresses B12-driven pro-inflammatory myeloid gene expression in stem, immature myeloid, and B cells in the bone marrow

To further investigate the origins of the B12-driven myeloid cell lineage differentiation bias, we performed scRNA-seq on donor CD45.2^+^ *Tet2*-deficient bone marrow cells isolated from control, B12 and butyrate treated host mice. Clusters were annotated according to Immgen gene expression profiles, Pseudotime analysis, and lineage surface marker expression (Fig.5A-C, Fig.S5A). The altered frequency of cells in several clusters by B12 treatment such as a reduction in B cells (cluster 0), and expansion of immature myeloid cells (clusters 6 and 11) were strikingly reverted upon butyrate co-supplementation (Fig.5D, Fig.S5B).

**Figure 5:**
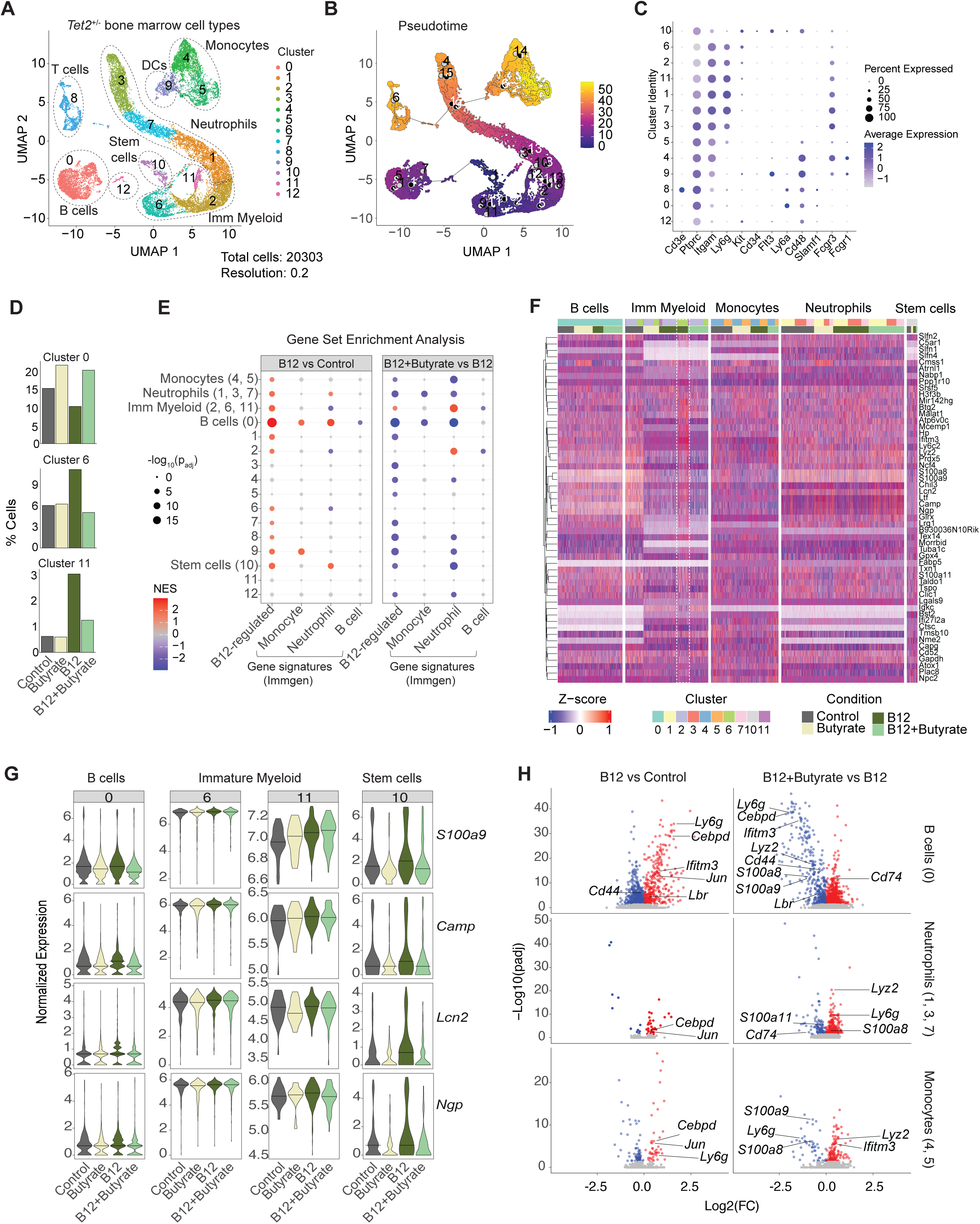
Butyrate suppresses B12-driven pro-inflammatory myeloid gene expression in *Tet2*-deficient stem cells, B cells, and immature myeloid cells in the bone marrow. (A) UMAP plot of scRNAseq classified using Immgen cell references with clustifyr and individual marker expression from cKit-enriched CD45.2^+^ *Tet2^+/−^* bone marrow cells isolated 8 months post-transplant from competitively reconstituted mice treated with Butyrate, high B12 or high B12+Butyrate. (B) Global pseudotime analysis of cell clusters across all cohorts. (C) Hematopoietic cell lineage and stem and progenitor hallmark gene expression grouped by cluster. (D) Relative percentages of cells in select clusters for each supplemented group. (E) Enrichment analysis by cluster or cell type (cluster group) of B12-upregulated genes identified from scRNAseq of splenic CD11b^+^ cells, and monocyte, neutrophil and B cell hallmark gene sets from Immgen. (F) Heatmap of splenic neutrophil and monocyte B12 high upregulated genes (see Figure 2) which show differential expression between B12+Butyrate vs B12 in the bone marrow, filtered for percent cell expression minimum. (G) Violin plots of normalized expression of representative B12-regulated genes *S100a9, Camp, Lcn2 and Ngp* across select clusters. (H) Volcano plots of differentially expressed genes in B cells (Cluster 0), neutrophils (clusters 1,3 and 7) and monocytes (clusters 4 and 5), with significantly (padj < 0.05) downregulated genes (blue) and upregulated (red) for the indicated comparisons. Select myeloid lineage and B12-regulated genes labeled.

We next performed a gene set enrichment analysis using B12-regulated genes identified from our splenic myeloid cell scRNA-seq analysis (Table S2) in addition to lineage-defining gene signatures of monocytes, neutrophils and B cells sourced from Immgen. Comparing B12-induced gene expression changes to control (IP supplementation vs standard diet alone) we observed a significant enrichment of B12-upregulated, monocyte and neutrophil signature genes most robustly in B cells (cluster 0), stem cells (cluster 10) and immature myeloid cells (clusters 2, 6 and 11 combined) – all of which were reverted upon butyrate co-treatment (Fig.5E). Many of these clusters showed the greatest number of differentially expressed genes (DEGs) when comparing B12 vs control and B12+Butyrate vs B12 treatment alone (Fig.S5C). Stem cells, immature myeloid and monocyte cluster cells also showed significant restoration of B12 down-regulated genes upon butyrate co-treatment (Fig.S5D). Importantly, both B cells (cluster 0) and stem cells (cluster 10) in the bone marrow showed significant differential expression of B cell genes and increased expression of monocyte and neutrophil genes upon B12 treatment that were reversed by butyrate (Fig.S5E). B12 upregulated genes annotated from CD11b^+^ splenic cells in response to high B12 treatment, including *S100a8*, *S100a9*, *Camp*, *Lcn2*, *Ngp*, *Lyz2* and *Wfdc21* also showed significant gene expression changes in immature myeloid cells, B cells and stem cells as well as in monocytes and neutrophils in the bone marrow upon high B12 treatment that were suppressed by butyrate co-treatment (Fig.5F-H, Fig.S5F). These data show that B12 drives a myeloid lineage differentiation bias that originates in the stem and immature myeloid progenitor cell compartment, reduces the expression of B lineage commitment genes in favor of myeloid gene expression in stem cells and developing B cells and, most importantly, is reversible upon butyrate co-treatment.

## Discussion

Elevated Vitamin B12 has been implicated in increased risk of mortality, cardiovascular disease, cancer (particularly myeloid malignancies), and most recently CH (*21–24, 44*). The source of excess B12 in individuals at risk is not clear. Elevated serum vitamin B12 levels can be caused by underlying clinical conditions such as liver disease, where hepatocyte damage leads to release of stored B12 into circulation, or myeloproliferative disorders such as chronic myeloid leukemia, where increased levels of haptocorrin-bound B12 are observed (*23, 44*). Renal dysfunction and solid tumors have also been associated with elevated B12 levels, potentially due to decreased clearance or ectopic production of B12-binding proteins (*22*). Moreover, supplement use with high-dose parenteral or oral B12 can elevate plasma levels (*44*).

A recent study reported that *Tet2* deficiency drives liver microbiome dysbiosis triggering liver damage and autoimmune hepatitis in mice (*45*). This could in principle cause a release of B12 stores and an elevation in B12 serum levels but was not assessed. *Tet2*-deficiency also increases intestinal permeability, leading to gut bacterial dissemination and translocation to the liver even under normal dietary B12 supplementation (*6*). Individuals with CH have an increased risk of developing chronic liver disease such as non-alcoholic steatohepatitis (NASH) (*46*). Studies using a dietary model of NASH confirmed that *Tet2*-deficient hematopoiesis causes more severe liver inflammation, and increases the expression of downstream inflammatory cytokines in *Tet2*-deficient macrophages to accelerate disease progression (*46*), suggesting that CH is causative of this increased risk. Intriguingly, we found that even on a standard chow diet, *Tet2*-deficiency alone reduced the abundance of SCFA-producing gut bacteria in mice as per high B12 supplementation, although whether this effect is secondary to liver inflammation and B12 release requires further investigation.

Elevated B12 serum levels have not been linked to liver dysbiosis, however, a recent large population-based study suggested that high B12 levels may also be causative in driving chronic liver disease and not just a consequence of liver damage. Individuals genetically predisposed to elevated serum B12 had an increased risk of non-alcoholic fatty liver disease (NAFLD) (*47*) which primarily encompasses non-alcoholic steatohepatitis (NASH), isolated hepatic steatosis, and cirrhosis (*48*). This relationship was bi-directional, as genetic liability to NAFLD was also positively associated with higher serum B12 concentrations (*47*). Taken together with our findings, these studies provide an additional layer of inflammatory-mediated positive feedback in *Tet2*-deficient CH progression that implicates microbial dysbiosis and liver damage as both a source and consequence of high B12 serum levels.

B12 may also exert cell-intrinsic metabolic changes in hematopoietic cells as described for other B vitamins, such as B6 and B2, that together with B12 regulate 1C metabolism involving folate (B9) and methionine recycling. Both B6 and B2 maintain the proliferation of AML cells by regulating metabolic enzymes essential for nucleotide and amino acid synthesis (*49, 50*) and antifolates like methotrexate were some of the first chemotherapeutic agents to show efficacy in the treatment of a variety of tumors by impairing nucleic acid synthesis (*51*). However, whether a detrimental effect of systemic B vitamin excess relates to their influence on gut microbial composition in the pathogenesis of blood cell malignancies has not been described. Recently, dietary vitamin D supplementation was shown to improve cancer immunotherapy through increasing the abundance and metabolic capacity of *Bacteroides* in the gut microbiome (*52*), highlighting the potential for micronutrients to influence microbial homeostasis and cancer treatment efficacy. We show that high B12 supplementation reduces beneficial commensals such as *Lactobacillus* and *Lachnospiraceae*, while enriching for *Bifidobacterium*, which have been linked to IL-17 and Th17-mediated inflammation (*53*). B12 also decreased the abundance of *Bacteroides*, SCFA-producing bacteria, and butanoate metabolism in the gut microbiome, and butyrate has previously been shown to decrease IL-17, IL-6, and IL-12 secretion from macrophages (*54–56*). Since SCFAs promote immune homeostasis, reduce cancer progression, and improve immunotherapy response (*57–59*), their depletion in the gut by excess B12 may exacerbate inflammation and CH progression.

Animal-derived foods provide the most abundant dietary source of B12 and are also high in protein. High protein consumption is known to cause a decrease in fecal SCFAs and alter microbial genera that resemble changes observed in patients with IBD (*11*). An unhealthy diet involving high meat consumption has also been linked to increased prevalence of CH (*20*). Increased meat consumption may elevate B12 and thus contribute to microbial dysbiosis and inflammation in predisposed individuals such as those carrying cancer-causing CH mutations in their blood cells.

*TET2* mutation is one of the most common drivers of CH and *Tet2*-deficient myeloid cells are known to be hyperresponsive to gut microbial signals (*6, 60*). We find that *Tet2*-deficient CH murine models supplemented with high B12 exhibited elevated levels of inflammatory cytokines associated with chronic inflammation in the gut (*61–63*), and progression of myeloid malignancy (*64, 65*): features already conferred by *TET2* deficiency but enhanced by high B12, suggesting a microenvironmental cooperation with TET2 loss of function. Additionally, B12 increased MHC class II expression in *Tet2*-deficient monocytes, a marker of poor prognosis in AML (*37*), and enhanced LPS-induced cytokine responses of myeloid cells *in vitro*, the latter implying a cell-intrinsic synergy between B12 and *Tet2-*deficiency. Structurally, B12 is a large cobalt-containing corrinoid with similarity to heme (*66*), which can activate TLR4 signaling and acute-phase responses (*67, 68*). Gene set enrichment analysis in our scRNAseq data confirmed that B12 treatment upregulated TLR4 signaling in monocytes and neutrophils. A recent study identified ADP-heptose, a biproduct of gram-negative bacteria, can induce inflammatory signaling responses that drive CH progression (*69*). Given that heme can amplify gram-negative derived LPS responses via MD-2/TLR4 (*70, 71*), our findings raise the possibility that B12, another microbially-derived metabolite, may directly engage similar innate immune pathways, to which *Tet2*-deficent cells are known to be hyperresponsive.

SCFAs, particularly butyrate and propionate, support intestinal homeostasis through anti-inflammatory, metabolic, and barrier-enhancing effects that have made them attractive therapeutic candidates for gut inflammatory disorders. SCFAs are produced through the microbial fermentation of dietary fiber and act as energy sources for colonocytes, while also exerting anti-inflammatory effects via inhibition of NF-κB signaling and suppression of pro-inflammatory cytokines such as IL-6, TNF-α, and IL-1β (*12, 15, 72*). Butyrate enhances epithelial barrier integrity by upregulating tight junction proteins and mucins, including MUC2, and by downregulating inflammasome activation, thereby preventing gut permeability and bacterial translocation (*12, 43, 54, 73*). IBD patients commonly exhibit gut dysbiosis characterized by the depletion of SCFA-producing bacteria, including *Faecalibacterium prausnitzii* and *Faecalibaculum rodentium*, which is thought to contribute to impaired mucosal healing and immune dysregulation (*6, 59*). In murine models, supplementation with butyrate or propionate restores microbial balance, limits intestinal inflammation, and improves epithelial integrity (*13, 42, 43*).

Butyrate may also exert its anti-inflammatory effects directly, by suppressing the expression of pro-inflammatory cytokines in intestinal macrophages via HDAC inhibition, leading to reduced colonic inflammation and colon cancer progression in mice (*55, 74*). The combined cell intrinsic and microenvironmental effects of a commensal metabolite such as butyrate are not only protective in IBD but have also been shown to attenuate graft-versus-host disease (*13*), and AML (*43*), by modulating barrier function. The therapeutic potential of SCFAs such as butyrate is further supported by their ability to promote the differentiation of regulatory T cells (Tregs), and suppress epithelial damage pathways, all of which converge to restore mucosal homeostasis and immune tolerance (*12, 13, 75, 76*).

Our findings support the development of SCFA-based therapies, such as butyrate supplementation, as non-toxic and mechanistically rational approaches to treating inflammation associated with CH and neoplastic conditions such as myeloid malignancies that have been linked to microbial dysbiosis and epithelial barrier dysfunction. Furthermore, the observation that B12 induces microbial shifts that impair SCFA production highlights a novel mechanism by which micronutrients influence pre-malignant hematopoiesis, suggesting that limiting excess B12 intake could be implemented as a preventative dietary intervention to mitigate CH-associated inflammation and risk of malignant transformation. Whether heightened inflammatory signaling is the underlying cause for increased disease risk and all-cause mortality in the presence of elevated serum B12 levels requires further investigation.

## Materials and Methods

### Mice

All animal protocols were approved by the University of Miami Institutional Animal Care Use committee. B6.SJL-Ptprc^a^Pepc^b^/BoyJ (CD45.1) and C57BL/6J (CD45.2) were purchased by Jackson Lab. Germline *Tet2*-deficient mice have been described previously (*29*) and are bred in a mouse facility at the University of Miami.

### Bone marrow competitive transplantation

Freshly dissected femur, tibia, and pelvis were isolated from donor *Tet2^+/+^* or *Tet2^+/−^* germline mice (CD45.2^+^) as well as *Tet2^+/+^* (CD45.1^+^) mice. Bones were flushed with sterile PBS using a 25-gauge needle and the isolated marrow was centrifuged for 5 minutes at 1500 RPM at room temperature. Cells were treated with ammonium-chloride-potassium (ACK) lysis buffer to lyse red blood cells. Cells were resuspended in PBS, passed through a 40 µm cell strainer and counted. Donor cells (0.5 x10^6^ per genotype, per mouse) were mixed 1:1 with support bone marrow (CD45.1^+^) and transplanted via retro-orbital injection into lethally irradiated female recipient mice. Mice were placed on respective treatments (Table S1) 1-month post-transplant.

Chimerism (CD45.2^+^ vs CD45.1^+^ cells) and B cell/T cell/Myeloid cell (BTM) frequencies (staining for B220, CD3, and CD11b, respectively) were monitored at 1-month intervals by flow cytometry of peripheral blood until 8 months post-transplant. Complete blood counts (CBCs) were acquired on a Heska Element HT5 hematology analyzer. At 8 months, mice were bled for plasma or serum collection, and fecal samples were collected before timed sacrifice. Chimerism and BTM differentials were assessed in the bone marrow, spleen, and livers of mice by flow cytometry.

### High B12 supplementation

For B12 IP supplementation, mice were injected twice weekly with PBS (B12 Low and control groups) or 0.29 µg B12 in PBS (B12 High group), equivalent to a 1mg clinical dose in a 70kg human. This 0.29 µg B12 dose is around one-tenth of the maximum dose reported in mice with no toxicity (*18*). For B12 dietary supplementation experiments, mice were placed on control or high B12 diets 1-month post-transplant (Table S1).

### Flow Cytometry

Single cell suspensions were prepared from peripheral blood, bone marrow, spleen, or liver, red blood cells were ACK-lysed, and remaining cells were suspended in 3% FPS in PBS. Cells were incubated with 1:100 mouse Fc block for at least 15 min, then surface marker antibodies were added for at least 30 min. For the BTM staining, cells were stained for CD45.1, CD45.2, B220, CD3, and CD11b at 1:500 concentration. For HSPC staining, cells were stained for lineage markers (B220, CD4, CD8, CD11b, Ter119, Gr1), CD45.1, CD45.2, cKit, Sca-1, CD150, and CD48, and myeloid progenitors were stained for FcgR (CD16/32) and CD34 expression at 1:100 concentration.

### Plasma and Serum Collection

Blood collected retro-orbitally from mice was centrifuged for 10 minutes at 2,000 x g at 4°C in EDTA-coated Eppendorf tubes (for plasma isolation) or allowed to clot undisturbed at room temperature for 30 minutes before centrifuging in non-treated tubes (for serum isolation). Supernatant was carefully collected, flash frozen, and stored at –80°C.

### B12 Measurement and IP kinetics

C57BL/6J mice were IP injected with 0.29 µg B12 in PBS once and plasma was collected at 0 hours, 1 hour, 6 hours, 72 hours and 1 week. B12 was measured by ELISA with the Novus Biologicals Mouse Vitamin B12 ELISA Kit (Colorimetric) (NBP2-59958). Plasma samples were diluted 1:25 in provided sample diluent, and the kit was followed per manufacturer protocol. Plates were read on a Synergy 2 plate reader (Biotek).

### Cytokine Analysis

Cytokines were measured at 8 months post-transplant (7 months of diet) with the Proteome Profiler Mouse Cytokine Array Kit (R&D system, ARY006). 50 µL of plasma or serum was used per sample, and the kit was followed according to manufacturer’s instructions. Image density was quantitatively calculated using HLImage++ software and was normalized to the corresponding reference and group.

### Microbiome Shotgun Sequencing

Fecal bacterial DNA extraction and microbiome shallow shotgun sequencing (3 million reads/sample) was performed by CosmosID. Briefly per CosmosID, DNA from samples was isolated using the QIAGEN DNeasy PowerSoil Pro Kit, according to the manufacturer’s protocol. Extracted DNA samples were quantified using Qubit Flex fluorometer and Qubit™ dsDNA HS Assay Kit (Thermofisher Scientific). DNA libraries were prepared using the xGen DNA Library Prep Kit (IDT) and xGen Normalase UDI Primers with total DNA input of 1.5ng. Genomic DNA was fragmented using a proportional amount of IDT xGEN fragmentation enzyme. DNA libraries were purified using AMpure magnetic Beads (Beckman Coulter) and eluted in QIAGEN EB buffer. DNA libraries were quantified using Qubit fluorometer and Qubit™ dsDNA HS Assay Kit. Libraries were then sequenced on an Illumina NovaSeq X Plus platform 2×150bp. Comparative analysis was conducted on the CosmosID hub (https://www.cosmosid.com/) to determine alpha diversity (Shannon and Simpsons Indices). Additionally, LEFSE analysis was calculated within each diet cohort and genotype with an Alpha of 0.05 and a LDA cutoff of 2. Specifically, the non-parametric factorial Kruskal-Wallis sum-rank was used to detect features with significant differential abundance with respect to species. Linear Discriminant Analysis is then used to estimate effect size of each feature and rank features accordingly.

### SCFA producer calculations

Calculations were made by summing the percent composition of each SCFA-producing genus (*72*) detected: acetate (*Akkermansia*, *Bifidobacterium*, *Prevotella*, *Ruminococcus*), propionate (*Akkermansia*, *Bacteroides, Eubacterium, Ruminococcus, Veillonella*) and butyrate (*Bacteroides, Clostridium, Eubacterium, Faecalibacterium, Fusobacterium*, *Roseburia*, *Spirochaeta*, *Thermotoga*).

### Metagenomic Profiling

Metagenomic analysis was completed using MG-RAST server as previously described (*77, 78*). The data were analyzed using the KO KEGG platform in MG-RAST with relative abundance of sequence per functional category is reported per sample. Log_2_(Fold-Change) was determined in comparisons and a Student’s T-test used to determine significance.

### Butyrate Supplementation

Competitive BM reconstituted mice were treated from 1-month post-transplant. For B12 supplementation, mice were IP injected twice weekly with PBS or B12 as described previously. For butyrate supplementation, mice were given water with control (100mM NaCl) or butyrate (100mM NaButyrate) *ad libitum*. Mice were treated in 4 groups: control (NaCl + PBS IP), butyrate (NaButyrate + PBS IP), high B12 (NaCl + B12 IP), high B12 + butyrate (NaButyrate + PBS IP). Chimerism (anti-CD45.1, anti-CD45.2) and B cell/T cell/Myeloid cell (BTM) frequencies (anti-B220, anti-CD3, and anti-CD11b) were monitored at 2-month intervals by flow cytometry of peripheral blood until 8 months post-transplant. Complete blood counts (CBCs) were acquired on a Heska Element HT5 hematology analyzer. At 8 months, mice were bled for serum collection before timed sacrifice. Additionally at 8 months, intestinal permeability was assessed with a FITC-Dextran assay. Chimerism of the hematopoietic compartments was assessed by flow cytometry.

### Single-cell RNA sequencing

Spleens harvested from *Tet2*^+/−^ competitive transplants from the B12 low (n=4) and B12 high (n=2) treatment groups were isolated into single cell suspension and ACK-lysed. CD45.2^+^CD11b+ cells were purified by flow cytometry, and samples within each treatment cohort were pooled. Sequencing libraries were prepared using 10X Chromium Single Cell 3’ V3 chemistry and assayed on a NovaSeq 6000. Quality control passing FASTQ reads were aligned to reference genome mm10, assembled into transcripts, and packaged into cell-indexed, gene-level matrices using cellranger count (v7.0.0). Downstream analysis was performed with seurat (v.4.0.2) using default function parameters, unless otherwise noted. Ribosomal transcripts were first purged, then samples merged to a single dataset, which was normalized and scaled using scTransform. Uniform Manifold Approximation and Projection (UMAP) reductions were created using RunPCA, FindVariableFeatures, FindNeighbors, FindClusters and RunUMAP, relying on the top 15 loadings for Principal Component Analysis of the top 3,000 variable features. Final clustering resolution of 0.2 was chosen based on clustree analysis. Cluster-defining markers were identified using FindAllMarkers. Cluster-marker heatmaps were rendered with dittoseq using the pheatmap wrapper. Clusters were annotated based on reference datasets (e.g. ImmGen references) using celldex (v.1.14.0) and/or known lineage-or state-defining markers. Datapoints (i.e. cells) were excluded if they had: >10% mitochondrial transcripts (dead/dying cells), >15,000 total transcripts or > 4,000 genes per cells (potential doublets, multiples), or <500 genes per cells (partial cells). Pseudotime analysis was performed using Monocle3. For differential gene expression analysis, clusters 0 and cluster 6 were combined (“Neutrophils”) and clusters 3 and 5 were combined (“Monocytes”) based on cluster classification. Next, the function “FindMarkers” from Seurat v5.1.0 R package was used to find differentially expressed genes based on B12 treatment (B12 High vs B12 Low). Wilcoxon Rank Sum test was used. Gene set enrichment analysis was performed on DEGs from the neutrophil clusters using Enrichr(*79*).

<u>Single-cell RNA-sequencing of the butyrate supplemented treatment cohort</u>, three viably frozen bone marrow samples per group were thawed, pooled separately per treatment-group, and flow cytometry sorted for live (DAPI^-^) CD45.2^+^ cells; samples were also sorted by cKit marker expression. After sorting, live CD45.2^+^ cells were pooled 50:50 cKit^+^:cKit^-^. Cytometry-sorted pooled cells were counted and evaluated for viability by brightfield and fluorescent microscopy by Nexcelom Cellometer K2 with ViaStain AOPI Staining Solution using the “Primary Tumor Dissociation” protocol. All sample viability was greater than 80% with low debris and aggregate frequency. 10,000 cells per sample were targeted for sequencing. Single cell isolation and library preparation was performed on the 10X Genomics Chromium platform using the Chromium Next GEM Single Cell 3’ Reagents Kit (v3.1). cDNA and gene expression libraries were evaluated for quality on an Agilent 5200 Fragment Analyzer. Libraries were sequenced on a NovaSeq X Plus using paired end, 150bp sequencing to a minimum 400M reads. Fastq files were barcode separated and aligned to the genome using cellranger multi (v7.1.0) and the GRCm39-2024-A reference from 10X Genomics. Cellranger output was read into a Seurat object. Ribosomal genes were filtered out. Cells were filtered to unique RNA features between 500-3500, RNA count below 20000, and mitochondrial genes below 15%. Expression was normalized by SCTransform with percent.mt regressed out. Nearest neighbor determination was made using FindNeighbors using dims 1:20 and “pca” reduction, followed by cluster determination using FindClusters, setting the resolution to 0.2. Uniform Manifold Approximation and Projection (UMAP) was performed using RunUMAP with the same dimensions and reduction as FindNeighbors. Pseudotime analysis was performed with Monocle3 (v1.4.26), setting cluster 10 as the root node. Differential marker detection was performed using the FindMarkers function and the MAST differential test (v1.32.0); each individual cluster or group of clusters defining a cell type was tested for each pairwise combination of treatments. Significant differentially expressed genes were defined as p_adj_ < 0.05. B12-regulated gene sets were created from the splenic high/low B12 Neutrophil and Monocyte differentially expressed genes (Supplementary Table 2). Immgen reference expression was retrieved using celldex (v1.12.0). Immgen gene sets were created by running findMarkers from scran (v1.34.0) on the log counts from the Immgen reference set using the wilcox rank sum test across all pairwise combinations of cell type. The top 100 (B cells) or 150 genes (Monocytes and Neutrophils) by summary AUC were used to define the gene set. GSEA was performed against the Immgen and B12 gene sets on the bone marrow Butyrate + B12 differential fold changes from FindMarker detection using fgsea (v1.32.2). Heatmap of B12-responsive genes was generated by filtering for significantly differential genes in the B12+Butyrate vs B12 comparison which overlapped with genes from the B12-upregulated gene set from the splenic dataset. Cell types of interest were separately z-score normalized, and color scale capped at +/−1. Heatmaps of Immgen cell-type gene signatures were generated using gene sets as described above, except using a summary AUC > 0.999 to generate a larger gene set, capping gene sets at 500. Gene sets were intersected with significant differentially expressed from the clusters indicated, from any pairwise comparison. Heatmaps were made with ComplexHeatmap (v2.22.0). All other plots were generated in R (v4.4.3) with Seurat (v5.3.0) or ggplot2 (v3.5.2).

### FITC-Dextran Assay

Food was removed from mouse cages for 4 hours prior to mice being gavaged with 150 µL of FITC-Dextran (3-5 kD @ 80 mg/mL). Mice were bled after one hour and plasma isolated as previously described. 100 µL serum was pipetted into a 96 well plate and fluorescence was measured with 485 nm excitation/535 emission on a Synergy 2 plate reader (Biotek). FITC-Dextran levels were calculated with a standard curve, and values were normalized to the mean of the standard diet group.

### Retroviral transduction

Raw 264.7 cells were transduced with retroviral constructs containing *shRenilla* (80) (*shRen)* or *shTet2* (29) (generated by Mirimus and previously described (*30*)) using Polybrene and pCL packaging vector. Cells were selected with puromycin at 1 µg/mL for 1 week.

### Raw 264.7 cell culture

*shRen* and *shTet2* Raw 264.7 cells were grown in a specially formulated RPMI without B12 (GIBCO). Cells were split into three B12 groups: Low B12 (0ng/L B12), Control (5 ng/L B12) and High B12 (10µg/L B12). Three biological replicates per B12 treatment were grown out to 4 months. For LPS experiments, cells were stimulated with LPS (1 µg/mL) overnight for 16 hours and harvested for RT-qPCR.

### RT-qPCR

Cells were homogenized with QIAshredders, and total RNA extracted using RNeasy Plus Mini Kit. For mRNA quantification, total RNA was used for cDNA synthesis using the High-Capacity RNA-to-cDNA Kit. Real-time PCR reactions were carried out using SYBR Select Master Mix on a QuantStudio 5 Real-Time PCR System (Applied Biosystems, A28575). Genes of interest were normalized against *Hprt,* and relative expression was determined using the 2^(-ΔΔCT)^ method. See Supplementary Table 3 for oligonucleotide sequences used.

### Quantification and Statistical Analysis

All p-values were calculated using a student’s t test GraphPad Prism 9 software. Analyses were not performed under specific randomization or blinding protocol. Statistically significant differences are indicated with asterisks in figures with the accompanying p value in the legend. Error bars in figures indicate standard deviation (SD) or standard error of the mean (SEM) for the number of replicates, as indicated in the figure legend.

## Acknowledgements

We thank the SCCC and Miller School of Medicine core facilities, including the Flow Cytometry, Oncogenomics, and Cancer modeling shared resources. Core facilities are supported by Sylvester Cancer Center Core support grants NCI SCCC-CCSG-P30 and 1P30CA240139-01. This research was also supported by the NIH NCI 1R01CA282453-01 (L Cimmino), Leukemia & Lymphoma Society (7028–22), and Castaways Against Cancer Foundation.

## Author Contributions

PL designed, performed, and analyzed *in vitro* and *in vivo* experiments, performed bioinformatics analysis and writing of the manuscript. TL performed mouse experiments, cell isolation, and analysis. JB performed bioinformatics analysis. KD, ML, and AP assisted with *in vivo* experiments. BS and VS assisted with colony maintenance. AA performed *in vitro* cell culture assays. LN, MD, and AV assisted with scRNAseq analysis. PS and SR contributed to the direction of the study. FB supervised bioinformatics analysis by JB. LC supervised the design, analysis, and interpretation of experiments, project management, and writing of the manuscript.

## Competing Interests

The authors declare they have no competing interests.

## Data Availability

Raw fecal shotgun sequencing datasets are available in the NCBI repository (SRA: PRJNA1122552). Raw and analyzed scRNAseq datasets are available at NCBI: GSE306040 (reviewer access token: yvofocearzeznij). Remaining data needed to support the conclusion of this manuscript are included in the main text and supplementary materials. Any additional information required to support the data reported in this paper is available from the lead contact upon request.

## Notes

### Competing Interest Statement

The authors have declared no competing interest.

