## Supplemental Figures and Legends for "B12 promotes gut dysbiosis and an inflammatory microenvironment that potentiates *Tet2*-deficient hematopoiesis"

#### Supplementary Figure Legends

**Figure S1. High B12 supplementation via intraperitoneal injection or increased dietary supplementation exacerbates *Tet2*-deficient myelopoiesis in mice.**

(A) Plasma B12 levels (pmol/L) measured by ELISA upon single IP injection of B12 (0.29µg in PBS).
(B) Red blood cell (RBC), hemoglobin (Hgb) and mean corpuscular volume (MCV) measurements in peripheral blood, 8 months after BM transplant with altered B12 supplementation.
(C) B, T and Myeloid cell frequencies measured by flow cytometry of $B220^+$, $CD3^+$ and $CD11b^+$ cells within the $CD45.2^+$ compartment in the peripheral blood (PB), bone marrow (BM), and spleen (SP) of mice at 8 months post-transplant.
(D) Schematic of $CD45.2^+$ $Tet2^{+/-}$ bone marrow (BM) mixed 1:1 with $CD45.1^+$ BM, transplanted into congenic mice placed on control or high B12 diet from 1-month post-transplant and monitored for 12 months.
(F-J) Hematopoietic phenotypes of competitively transplanted dietary B12 mice at 12 months post-transplant.
(E) Total WBC counts (k/µL)
(F) Total frequency of $CD45.2^+$ in peripheral blood (PB) bone marrow (BM) and spleen (SP) and $CD45.2^+$ B, T, and Myeloid cell frequencies in the BM and SP
(G) Myeloid to B cell (M/B) ratio measured by flow cytometry of $CD11b^+$ vs $B220^+$ cells in PB, BM and SP
(H) Frequency of $CD45.2^+$ Lineage negative (Lin-) $cKit^+$ (LK) cells, lineage negative $cKit^+$ $Sca1^+$ (LSK) cells and LSK subsets that are $CD150^+CD48^-$ (HSCs) and $CD150^-CD48^+$ (myeloid primed multipotent progenitors) within the BM compartment
(I) Frequency of common myeloid progenitor (CMP), megakaryocyte and erythroid progenitor (MEP) and granulocyte and macrophage progenitor (GMP) cells in the $CD45.2^+$ LK compartment

Panels show mean and STD of n = 3-5 mice per group, $^*p < 0.05$, $^{**}p < 0.005$, $^{***}p < 0.0005$.

**Figure S2: High B12 supplementation increases inflammatory gene expression in *Tet2*-deficient myeloid cells.**

(A) Heatmap of relative expression of the top 10 defining genes per cluster.
(B) Violin plots of normalized expression of *S100a9* across all scRNAseq clusters.
(C) Venn Diagrams of overlapping DEGs in the Monocyte and Neutrophil clusters.
(D) Enriched gene sets in commonly up- and downregulated DEGs in the Neutrophil and Monocyte clusters from the KEGG Pathway Database. Significance cutoff of padj < 0.05.
(E) Enriched gene sets in commonly up- and downregulated DEGs in the Neutrophil and Monocyte clusters from the Reactome Pathway Database. Significance cutoff of padj < 0.05.

**Figure S3: *Tet2*-deficient hematopoiesis alters the gut microbiome leading to reduced SCFA butyrate-producing bacteria.**

(A) Schematic of $CD45.2^+$ $Tet2^{+/+}$ or $Tet2^{+/-}$ bone marrow (BM) mixed 1:1 with $CD45.1^+$ BM transplanted into congenic mice monitored for 8 months with fecal pellets collected.
(B) $CD45.2^+$ percentage of peripheral blood by flow cytometry.
(C-F) Hematopoietic phenotypes of competitively transplanted mice at month 8 post-transplant.
(C) Total white blood cells (WBC)
(D) Percent $CD45.2^+$ cells in the bone marrow (BM) and spleen (SP)
(E) M/B ratio measured by flow cytometry of $CD11b^+$ vs $B220^+$ cells within the $CD45.2^+$ compartment in the bone marrow (BM) and spleen (SP)
(F) Percent $CD45.2^+$ cells within the $CD11b^+$ myeloid compartment in the liver
(G) Alpha diversity levels measured by Shannon and Simpson's indices in fecal pellets from $Tet2^{+/+}$ and $Tet2^{+/-}$ competitive transplant mice.
(H) Microbiome composition at the phylum level represented as individual mice and averaged per group in fecal pellets from $Tet2^{+/+}$ and $Tet2^{+/-}$ competitive transplant mice.
(I) Composition levels of acetate, propionate, and butyrate-producing genera in the fecal microbiome of $Tet2^{+/+}$ and $Tet2^{+/-}$ competitive transplant mice.
(J) Bar graphs of linear discriminant analysis (LDA) score calculated by linear discriminant analysis effect size (LEFSE) for species in the fecal microbiome of $Tet2^{+/+}$ and $Tet2^{+/-}$ hosts .
(K) Dot plot of enriched KEGG pathways from metagenomic analysis comparing the fecal microbiomes of $Tet2^{+/+}$ and $Tet2^{+/-}$ host mice using the MG-RAST pipeline.

Panels show mean and STD of n=8-9 mice per group, $*p < 0.05$, $**p < 0.005$, $***p < 0.0005$.

**Figure S4: Butyrate treatment ameliorates the B12-driven myeloid lineage bias of both $Tet2^{+/-}$ and $Tet2^{+/+}$ hematopoiesis.**

(A-C) Hematopoietic phenotypes at 8 months post-transplant.
(A) Bar graph of red blood cell (RBC), hemoglobin (Hgb) and mean corpuscular volume (MCV) in peripheral blood.
(B) B, T and Myeloid cell frequencies measured by flow cytometry of $B220^+$, $CD3^+$ and $CD11b^+$ cells within the $CD45.2^+$ compartment in the peripheral blood (PB), bone marrow (BM), and spleen (SP).
(C) M/B ratio of $CD45.1^+$ wild-type support cells measured by flow cytometry of $CD11b^+$ vs $B220^+$ cells in PB, BM and SP.
(D) Relative FITC fluorescence measured in plasma of mice gavaged with FITC-dextran at month 8 prior to sacrifice of supplemented mice.

Panels show mean and STD of n = 5-8 mice per group, $*p < 0.05$, $**p < 0.005$, $****p<0.00005$.

**Figure S5: Butyrate suppresses aberrant B12-driven myeloid gene expression in *Tet2*-deficient stem cells and B cells in the bone marrow.**

(A) UMAP plots of scRNAseq classified using Immgen cell references with clustifyr and individual marker expression from cKit-enriched CD45.2$^{+}$ *Tet2*$^{+/-}$ bone marrow cells isolated 8 months post-transplant from competitively reconstituted mice treated with Butyrate, high B12 or high B12+Butyrate compared to Control.
(B) Relative percentages of cells in all clusters for each supplemented group. Dashed boxes signify B12-modulated, Butyrate-responsive clusters.
(C) Number of significantly differentially expressed genes per cluster/cell type for the indicated differential comparisons.
(D) Enrichment analysis by cluster of B12-downregulated genes identified from scRNAseq of splenic CD11b$^{+}$ cells.
(E) Heatmaps of B cell, monocyte and neutrophil signature gene sets derived from Immgen cell references which are differentially regulated in B cells (cluster 0) and Stem Cells (cluster 10) from mice in the indicated treatment groups.
(F) Violin plots of normalized expression of additional splenic CD11b$^{+}$ B12-upregulated genes (*S100a8, Lyz2* and *Wfdc21*) in BM B cells (cluster 0) and Stem Cells (cluster 10) from mice in the indicated treatment groups.

### Figure S1

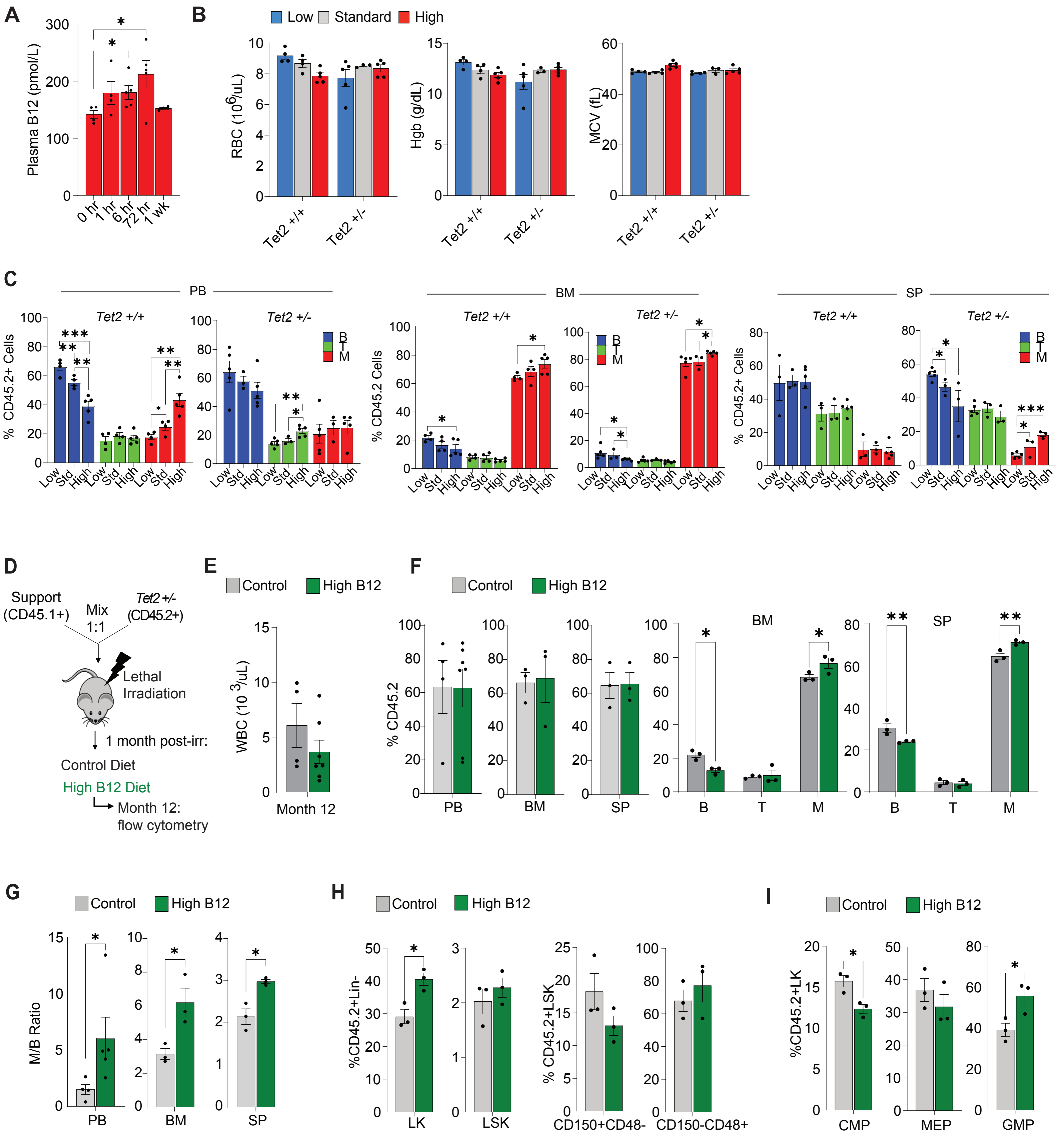

A
Plasma B12 (pmol/L)
0 hr
1 hr
6 hr
72 hr
1 wk
B
Low
Standard
High
RBC (10^6/uL)
Hgb (g/dL)
MCV (fL)
Tet2 +/+
Tet2 +/-
C
PB
BM
SP
Tet2 +/+
Tet2 +/-
% CD45.2+ Cells
% CD45.2 Cells
B
T
M
Low
Std
High
D
Support (CD45.1+)
Mix 1:1
Tet2 +/- (CD45.2+)
Lethal Irradiation
1 month post-irr:
Control Diet
High B12 Diet
Month 12: flow cytometry
E
Control
High B12
WBC (10 3/uL)
Month 12
F
% CD45.2
PB
BM
SP
G
M/B Ratio
H
%CD45.2+Lin-
LK
LSK
% CD45.2+LSK
CD150+CD48-
CD150-CD48+
I
%CD45.2+LK
CMP
MEP
GMP

#### Figure S2

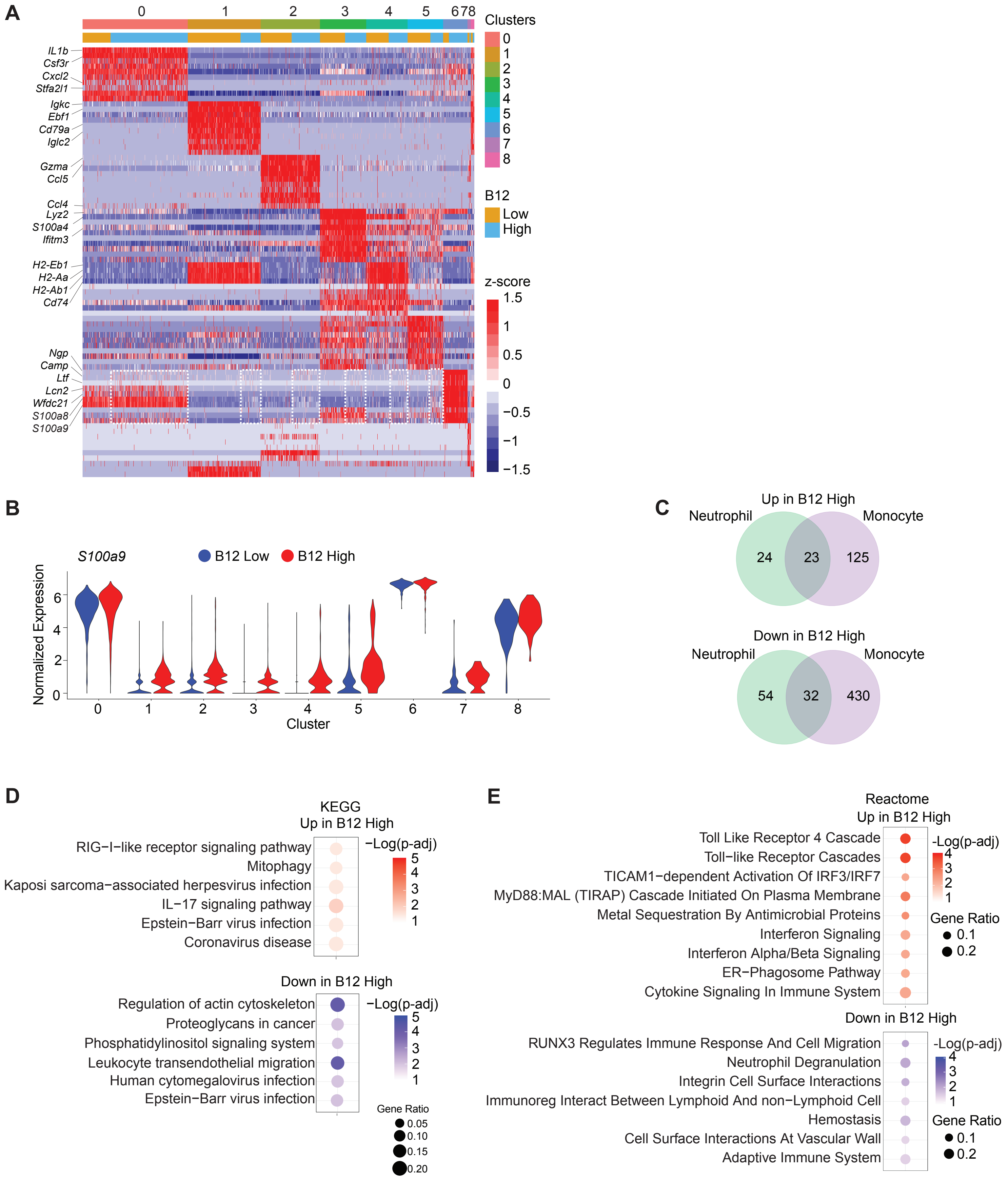

A
0 1 2 3 4 5 6 7 8
Clusters
0
1
2
3
4
5
6
7
8
B12
Low
High
z-score
1.5
1
0.5
0
−0.5
−1
−1.5
IL1b
Csf3r
Cxcl2
Stfa2l1
Igkc
Ebf1
Cd79a
Iglc2
Gzma
Ccl5
Ccl4
Lyz2
S100a4
Ifitm3
H2-Eb1
H2-Aa
H2-Ab1
Cd74
Ngp
Camp
Ltf
Lcn2
Wfdc21
S100a8
S100a9
B
S100a9
B12 Low
B12 High
Normalized Expression
Cluster
C
Up in B12 High
Neutrophil
Monocyte
24
23
125
Down in B12 High
Neutrophil
Monocyte
54
32
430
D
KEGG
Up in B12 High
RIG−I−like receptor signaling pathway
Mitophagy
Kaposi sarcoma−associated herpesvirus infection
IL−17 signaling pathway
Epstein−Barr virus infection
Coronavirus disease
−Log(p-adj)
Down in B12 High
Regulation of actin cytoskeleton
Proteoglycans in cancer
Phosphatidylinositol signaling system
Leukocyte transendothelial migration
Human cytomegalovirus infection
Epstein−Barr virus infection
−Log(p-adj)
Gene Ratio
0.05
0.10
0.15
0.20
E
Reactome
Up in B12 High
Toll Like Receptor 4 Cascade
Toll−like Receptor Cascades
TICAM1−dependent Activation Of IRF3/IRF7
MyD88:MAL (TIRAP) Cascade Initiated On Plasma Membrane
Metal Sequestration By Antimicrobial Proteins
Interferon Signaling
Interferon Alpha/Beta Signaling
ER−Phagosome Pathway
Cytokine Signaling In Immune System
-Log(p-adj)
Gene Ratio
0.1
0.2
Down in B12 High
RUNX3 Regulates Immune Response And Cell Migration
Neutrophil Degranulation
Integrin Cell Surface Interactions
Immunoreg Interact Between Lymphoid And non−Lymphoid Cell
Hemostasis
Cell Surface Interactions At Vascular Wall
Adaptive Immune System
-Log(p-adj)
Gene Ratio
0.1
0.2

### Figure S3

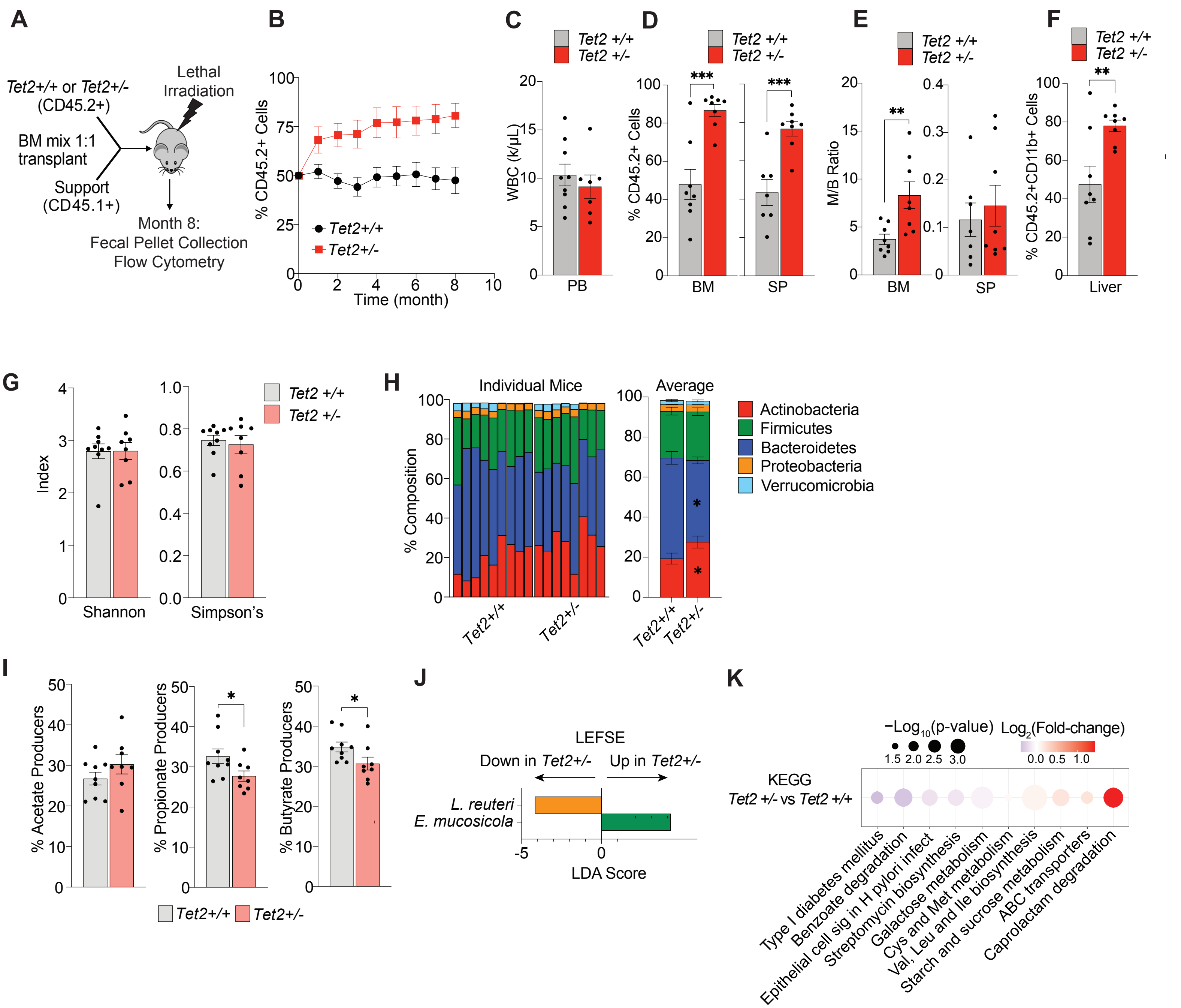
A
Tet2+/+ or Tet2+/-
(CD45.2+)
Lethal
Irradiation
BM mix 1:1
transplant
Support
(CD45.1+)
Month 8:
Fecal Pellet Collection
Flow Cytometry
B
% CD45.2+ Cells
Time (month)
Tet2+/+
Tet2+/-
C
Tet2 +/+
Tet2 +/-
WBC (k/μL)
PB
D
% CD45.2+ Cells
BM
SP
E
M/B Ratio
BM
SP
F
% CD45.2+CD11b+ Cells
Liver
G
Index
Shannon
Simpson's
H
Individual Mice
Average
% Composition
Actinobacteria
Firmicutes
Bacteroidetes
Proteobacteria
Verrucomicrobia
I
% Acetate Producers
% Propionate Producers
% Butyrate Producers
J
LEFSE
Down in Tet2+/-
Up in Tet2+/-
L. reuteri
E. mucosicola
LDA Score
K
−Log10(p-value)
Log2(Fold-change)
KEGG
Tet2 +/- vs Tet2 +/+
Type I diabetes mellitus
Benzoate degradation
Epithelial cell sig in H pylori infect
Streptomycin biosynthesis
Galactose metabolism
Cys and Met metabolism
Val, Leu and Ile biosynthesis
Starch and sucrose metabolism
ABC transporters
Caprolactam degradation

#### Figure S4

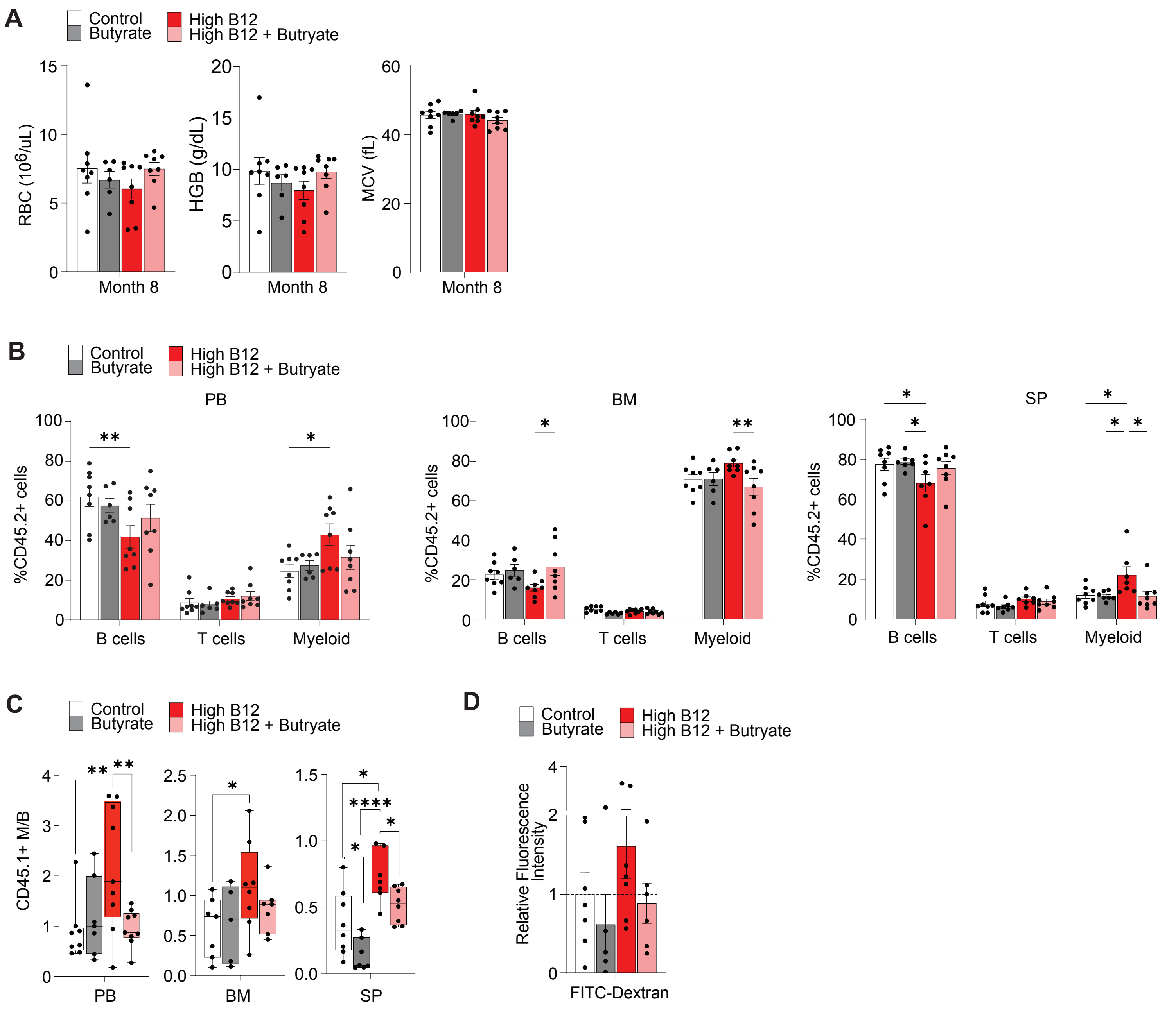

A
Control
Butyrate
High B12
High B12 + Butryate
RBC (10^6/uL)
HGB (g/dL)
MCV (fL)
Month 8
B
PB
BM
SP
%CD45.2+ cells
B cells
T cells
Myeloid
C
CD45.1+ M/B
PB
BM
SP
D
Relative Fluorescence Intensity
FITC-Dextran

### Figure S5

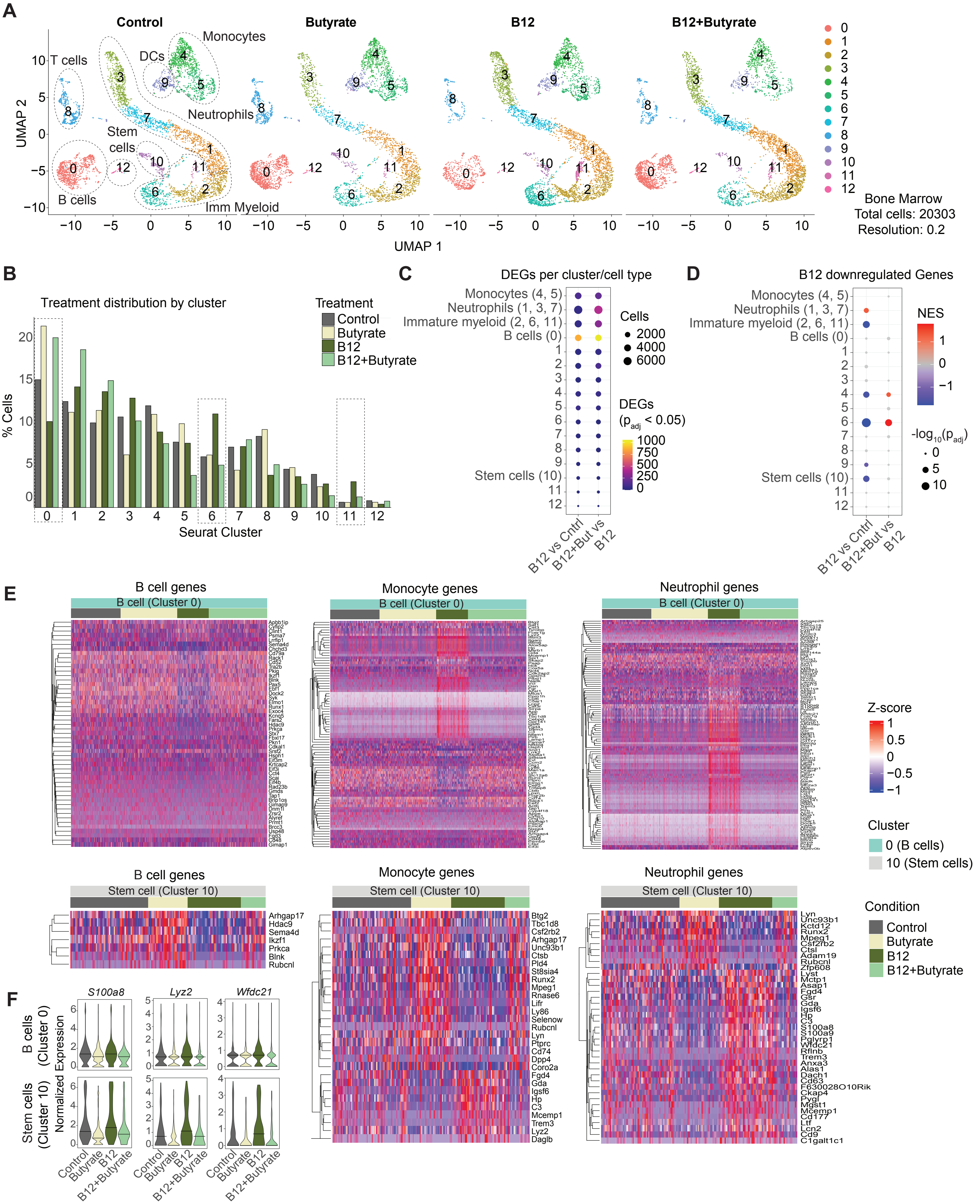

A
Control
Butyrate
B12
B12+Butyrate
T cells
DCs
Monocytes
Neutrophils
Stem cells
B cells
Imm Myeloid
UMAP 2
UMAP 1
0
1
2
3
4
5
6
7
8
9
10
11
12
Bone Marrow
Total cells: 20303
Resolution: 0.2
B
Treatment distribution by cluster
% Cells
Seurat Cluster
Treatment
Control
Butyrate
B12
B12+Butyrate
C
DEGs per cluster/cell type
Monocytes (4, 5)
Neutrophils (1, 3, 7)
Immature myeloid (2, 6, 11)
B cells (0)
Stem cells (10)
Cells
2000
4000
6000
DEGs ($p_{adj}$ < 0.05)
1000
750
500
250
0
B12 vs Cntrl
B12+But vs B12
D
B12 downregulated Genes
NES
$-log_{10}(p_{adj})$
B12 vs Cntrl
B12+But vs B12
E
B cell genes
Monocyte genes
Neutrophil genes
B cell (Cluster 0)
Stem cell (Cluster 10)
Z-score
Cluster
0 (B cells)
10 (Stem cells)
Condition
Control
Butyrate
B12
B12+Butyrate
Arhgap17
Hdac9
Sema4d
Ikzf1
Prkca
Blnk
Rubcnl
Btg2
Tbc1d8
Csf2rb2
Arhgap17
Unc93b1
Ctsb
Pld4
St8sia4
Runx2
Mpeg1
Rnase6
Lifr
Ly86
Selenow
Rubcnl
Lyn
Ptprc
Cd74
Dpp4
Coro2a
Fgd4
Gda
Igsf6
Hp
C3
Mcemp1
Trem3
Lyz2
Daglb
Lyn
Unc93b1
Kctd12
Runx2
Mpeg1
Csf2rb2
Ctsl
Adam19
Rubcnl
Zfp608
Lyst
Mctp1
Asap1
Fgd4
Gsr
Gda
Igsf6
Hp
C3
S100a8
S100a9
Pglyrp1
Wfdc21
Rflnb
Trem3
Anxa3
Alas1
Dach1
Cd63
F630028O10Rik
Ckap4
Pygl
Mgst1
Mcemp1
Cd177
Ltf
Lcn2
Cd9
C1galt1c1
F
S100a8
Lyz2
Wfdc21
B cells (Cluster 0)
Stem cells (Cluster 10)
Normalized Expression
Control
Butyrate
B12
B12+Butyrate
